# *RUNX1* isoform disequilibrium in the development of trisomy 21 associated myeloid leukemia

**DOI:** 10.1101/2022.03.07.483334

**Authors:** Sofia Gialesaki, Daniela Bräuer-Hartmann, Raj Bhayadia, Oriol Alejo-Valle, Hasan Issa, Enikő Regenyi, Christoph Beyer, Maurice Labuhn, Christian Ihling, Andrea Sinz, Markus Glaß, Stefan Hüttelmaier, Sören Matzk, Lena Schmid, Farina Josepha Strüwe, Sofie-Katrin Kadel, Dirk Reinhardt, Marie-Laure Yaspo, Dirk Heckl, Jan-Henning Klusmann

## Abstract

Aneuploidy is a hallmark of cancer, but its complex nature limits our understanding of how it drives oncogenesis. Gain of chromosome 21 (Hsa21) is among the most frequent aneuploidies in leukemia and is associated with markedly increased leukemia risk. Here, we propose that disequilibrium of the *RUNX1* isoforms is key to the pathogenesis of trisomy 21 (i.e. Down syndrome)-associated myeloid leukemia (ML-DS). Hsa21-focused CRISPR-Cas9 screens uncovered a strong and specific *RUNX1* dependency in ML-DS. Mechanistic studies revealed that excess of RUNX1A isoform – as seen in ML-DS patients – synergized with the pathognomonic *Gata1s* mutation in leukemogenesis by displacing RUNX1C from its endogenous binding sites and inducing oncogenic programs in complex with the MYC cofactor MAX. These effects were reversed by restoring the RUNX1A:RUNX1C equilibrium or pharmacological interference with MYC:MAX dimerization. Our study highlights the importance of alternative splicing in leukemogenesis, even on a background of aneuploidies, and opens new avenues for developing specific and targeted therapies.

**Graphical Abstract:** 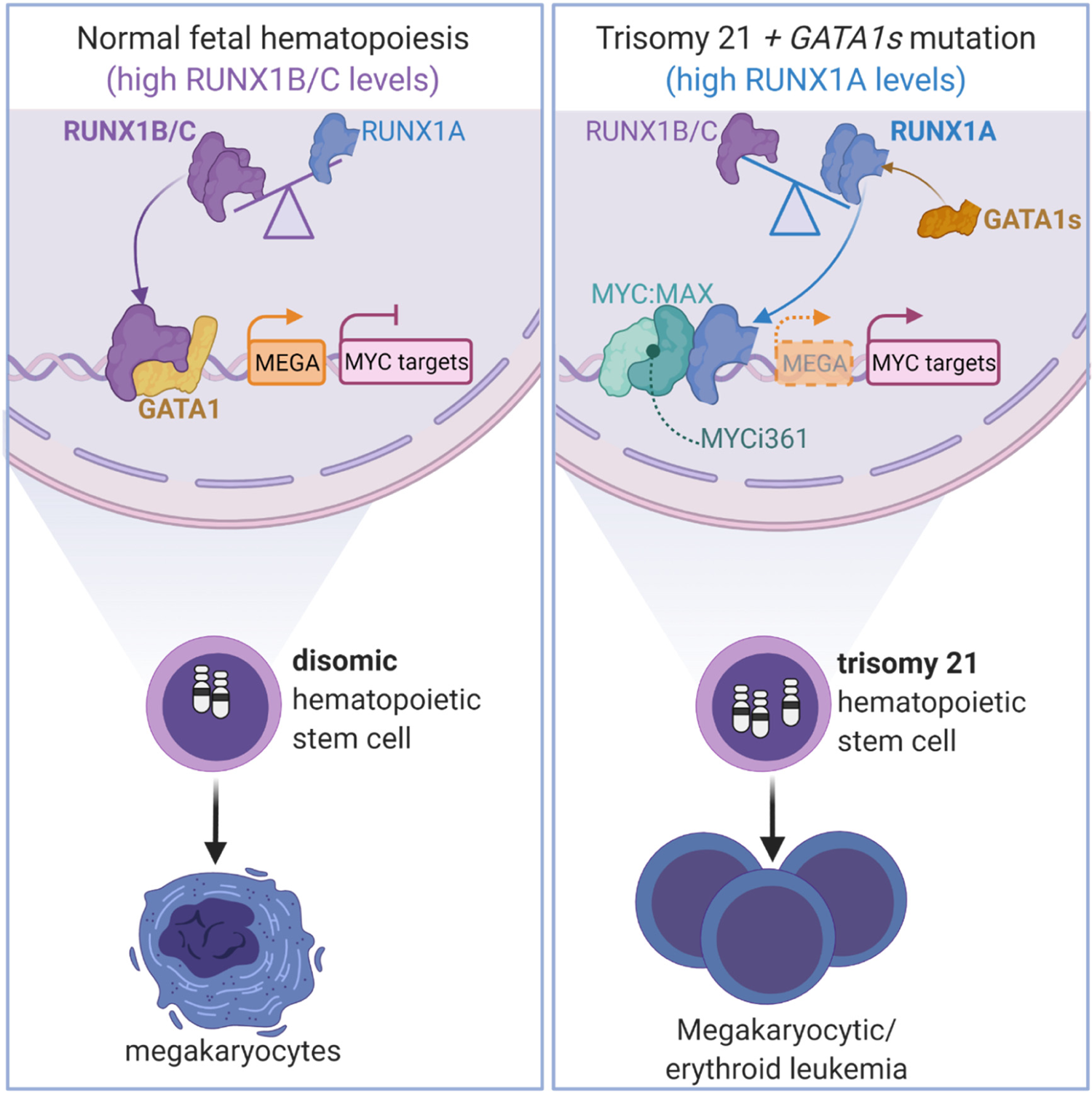

## Introduction

Major efforts in omics have mapped the cytogenetic, mutational and epigenetic landscape of cancer, including acute myeloid leukemia (AML) (Bolouri et al., 2018; Cancer Genome Atlas Research et al., 2013). Subsequent functional studies involving disease modeling in mice or human cells have pointed towards complex cooperation between common fusion oncogenes and recurrently mutated genes during disease initiation and progression. However, the contribution of aneuploidy to oncogenesis remains poorly understood (Ben-David and Amon, 2020), due to context-dependent effects, technical challenges, and a lack of appropriate models. Chromosome 21 (Hsa21) gain – one of the most frequent numerical alterations in leukemia (Duijf et al., 2013; Mitelman et al., 1990) – is no exception.

Myeloid leukemia associated with Down syndrome (ML-DS) and its preleukemic predecessor, transient abnormal hematopoiesis (TAM), are excellent paradigms for studying leukemic progression associated with trisomy 21. TAM is caused by a single genetic mutation in the transcription factor GATA1 in trisomic fetal stem and/or progenitor cells, which causes the exclusive expression of a shorter isoform known as GATA1s (Wechsler et al., 2002). We and others have shed light on the additional mutational events that are required for progression to overt ML-DS (Labuhn et al., 2019; Yoshida et al., 2013), however, far less is known about the role of trisomy 21 in creating the TAM/ML-DS disease phenotype. A critical region has been identified on Hsa21 that mediates the expansion of early hematopoietic progenitors observed in DS patients (Banno et al., 2016; Gurbuxani et al., 2004; Roy et al., 2012) and several Hsa21 genes – such as *ERG* (Birger et al., 2013; Ng et al., 2010), *DYRK1A* (Malinge et al., 2012), *CHAF1B* (Volk et al., 2018) and miR-125b (Alejo-Valle et al., 2021; Wagenblast et al., 2021) – have been postulated to play a role in leukemogenesis. On the other hand, Ts65dn mice that are trisomic for 104 orthologs of Hsa21 genes do not fully recapitulate the human phenotype in association with *GATA1s* (Klusmann et al., 2010), and the postulated factors are either located outside of the critical region, not overexpressed in trisomic fetal progenitor cells (Roy *et al*., 2012), or lack full leukemic potential in humans when combined with mutated *GATA1s* (Chou et al., 2012; Grimm et al., 2021; Malinge *et al*., 2012).

The transcription factor RUNX1, which is essential for the establishment of definitive hematopoiesis (North et al., 2002; Okuda et al., 1996; Wang et al., 1996), has attracted considerable attention as a candidate Hsa21 oncogene. While RUNX1 has been extensively studied in leukemia development (Ichikawa et al., 2013; Lam and Zhang, 2012), its expression is reduced in trisomic fetal progenitor cells (Roy *et al*., 2012) and its trisomy is dispensable for myeloproliferative disease in elderly Ts65dn mice (Kirsammer et al., 2008). Studies to date have not accounted for the fact that *RUNX1* is transcribed from two distinct promoters and undergoes alternative splicing, giving rise to three isoforms with diverse effects on hematopoiesis (Ghozi et al., 1996): *RUNX1C* is the most abundant isoform in definitive hematopoiesis, and *RUNX1A* and *RUNX1B* are differentially expressed throughout hematopoietic differentiation (Challen and Goodell, 2010; Fujita et al., 2001; Sroczynska et al., 2009). Interestingly, mice do not express *RUNX1A* – the short *RUNX1* isoform that lacks the transactivation domain – underlining important species-specific differences (Komeno et al., 2014), and contributing in part to the incomplete understanding of isoform-specific roles for RUNX1.

In this study, we systematically investigated protein coding genes of Hsa21 for dependency in ML-DS, and found that disequilibrium of the *RUNX1* isoforms – specifically, excess *RUNX1A –* is key to trisomy 21-associated leukemogenesis.

## Results

### CRISPR-Cas9 screen reveals RUNX1 dependency in ML-DS

In order to identify potential oncogenes on Hsa21 that contribute to the pathogenesis of TAM and ML-DS, we generated a lentiviral CRISPR-Cas9 library (1090 sgRNA) targeting the 218 annotated coding genes on Hsa21 and performed a dropout screen in the ML-DS cell line CMK, as well as in the erythroleukemia cell line K562 (**Figure 1A**). We identified 19 genes specifically required for the survival of CMK cells (**Figure 1B-C** and **Table S1**). Interestingly, we observed a strong depletion of *RUNX1*-targeting sgRNAs in CMK but not K562 cells (**Figure 1B-C**). Subsequent flow cytometry-based depletion assays using individual sgRNAs found *RUNX1* to be the most specific ML-DS dependency in our screen (**Figure 1C** and **Figure S1A-B**). These results were recapitulated in stable Cas9-expressing ML-DS patient-derived blasts *in vitro* and in fluorescence-based competitive transplantation assays *in vivo* (**Figure 1D-E** and **Figure S1C-D)**.

**Figure 1.**
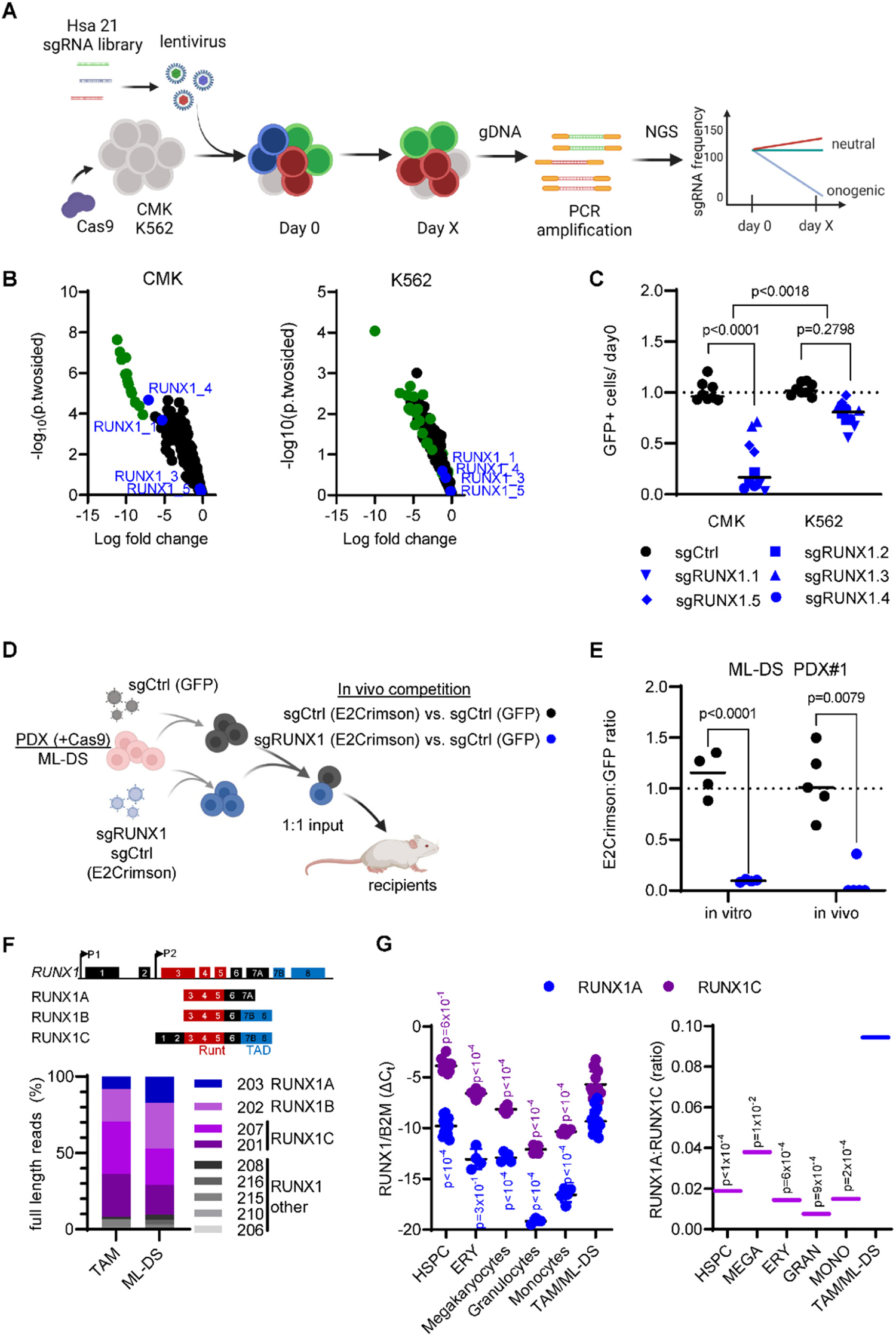
CRISPR-Cas9 screen reveals *RUNX1* dependency in ML-DS. (A) Schema of the high-throughput Hsa21 CRISPR-Cas9 *in vitro* screen. (B) Dot plots showing the log_10_ fold change and -log_10_ (p two-sided) of the Hsa21 sgRNAs in CMK (left) and K562 (right) cell lines. Green dots represent highly depleting sgRNAs in CMK cells that also deplete in K562s. Blue dots represent sgRNAs targeting *RUNX1*. (C) Dot plot showing the number of sgRNA-transduced (sgCtrl and sgRUNX1.1-1.5 indicated by shape) CMK and K562 cells (with stable Cas9 expression) after 14 days of culture normalized to day 0 (n=2 per sgRNA, two-tailed unpaired t-test). (D) Experimental setup for evaluating *RUNX1* knockout *in vivo*. ML-DS blasts (with stable Cas9 expression) were transduced with sgRUNX1 (E2Crimson^+^, sgRUNX1.1) or a sgCtrl (E2Crimson^+^) and mixed 1:1 with sgCtrl-transduced blasts (GFP^+^), before transplantation into sub-lethally irradiated recipient mice. (E) Dot plot showing the ratio of E2Crimson^+^ to GFP^+^ cells after 12 days of culture normalized to day 0 (left; n=4, two-tailed unpaired t-test) and in the bone marrow of mice sacrificed 6-8 weeks after transplantation (right; n=5, Mann Whitney test). (F) Schematic representation of the human RUNX1 genomic locus (upper panel) and the 3 main RUNX1 transcript isoforms (lower panel; not to scale). Functional exons encoding the DNA-binding domain (Runt; red) and transactivation domain (TAD; blue) are indicated. Bar graph showing RUNX1 isoform distribution (Oxford Nanopore sequencing) in polyA^+^-enriched RNA samples from TAM and ML-DS samples. (G) Expression of *RUNX1A* and *RUNX1C* isoforms normalized to the expression of β2-microglobulin (*B2M*) (left graph; two-way ANOVA) and *RUNX1A* to *RUNX1C* ratios (right graph; one-way ANOVA) in CD34^+^ HSPCs, erythrocytes, megakaryocytes, granulocytes and monocytes isolated from healthy donors, as well as in leukemic blasts from TAM/ML-DS patients. See also **Figure S1** and **Table S1-2**.

Considering that former studies have disputed the role of *RUNX1* in TAM/ML-DS pathogenesis (Bourquin et al., 2006; Kirsammer *et al*., 2008; Roy *et al*., 2012), we wondered whether deregulation of the *RUNX1* isoforms may underlie its dependency phenotype in ML-DS cells, rather than altered overall expression. Nanopore full length RNA sequencing revealed that the main isoforms – *RUNX1A*, *RUNX1B* and *RUNX1C* – are predominant in TAM and ML-DS (**Figure 1F** and **Table S2).** We thus further quantified the expression of these 3 isoforms in healthy hematopoietic cells (HSPCs, erythrocytes, megakaryocytes, granulocytes and monocytes) and TAM/ ML-DS blasts (**Figure 1G and Figure S1E**). *RUNX1C* is the predominant isoform in all cell types, but ML-DS samples presented with elevated expression of the *RUNX1A* isoform, resulting in a significantly higher *RUNX1A*:*RUNX1B/C* ratio compared to HSPCs or terminally differentiated cells (**Figure 1G** and **Figure S1F**). Notably, trisomy 21 is associated with an elevated *RUNX1A*:*RUNX1B/C* ratio, as determined by comparing fetal trisomy 21 and non-trisomic CD34^+^ HSPCs (**Figure S1E-F**). Moreover, GATA1s’ ability to regulate *RUNX1A* appears to be impaired compared to full-length GATA1. It shows reduced occupancy at the RUNX1 P1 und P2 promoter regions in *Gata1s*-FLCs and fails to repress *RUNX1A* in CRISPR-Cas9 edited human megakaryocytic/ erythroid precursors cultured *in vitro* (**Figure S1G-I**), suggesting a mechanism whereby trisomy 21 and GATA1s can synergize to amplify *RUNX1* isoform imbalance. Thus, our CRISPR-Cas9 screen of Hsa21 genes and subsequent *RUNX1* isoform expression analyses in patient samples suggest that *RUNX1* isoform disequilibrium and *RUNX1A* bias contribute to the pathogenesis of TAM and ML-DS.

### Increased RUNX1A:RUNX1C ratio induces malignant ML-DS phenotype

To probe the functional relevance of unbalanced *RUNX1* isoform expression in TAM and ML-DS development, we modulated the *RUNX1A*:*RUNX1C* ratio in neonatal CD34^+^ HSPCs of human origin and in patient-derived ML-DS blasts. Methylcellulose-based CFU assays showed the sustained growth of *RUNX1A*-expressing CD34^+^ HSPCs, as evidenced by increased replating capacity (**Figure 2A**). In culture conditions promoting megakaryocytic differentiation, *RUNX1C* induced terminal differentiation (CD41/61^+^CD42b^+^ cells), while *RUNX1A* expression induced the expansion of immature megakaryocytic cells and blocked their differentiation (**Figure 2B** and **Figure S2A-B**) – effects that were confirmed in collagen-based megakaryocytic colony-forming (CFU-Mk) assays (**Figure S2C**). To better understand the effects of *RUNX1A* and *RUNX1C* on lineage fate decision, we additionally cultured CD34^+^ HSPCs in conditions allowing for differentiation along the erythroid as well as megakaryocytic lineages. Whereas *RUNX1C* conferred a lineage bias towards megakaryopoiesis (CD41^+^/CD235^-^) over erythropoiesis (CD41^-^/CD235^+^), *RUNX1A* blocked differentiation along both lineages (**Figure 2C** and **Figure S2D-E**).

**Figure 2.**
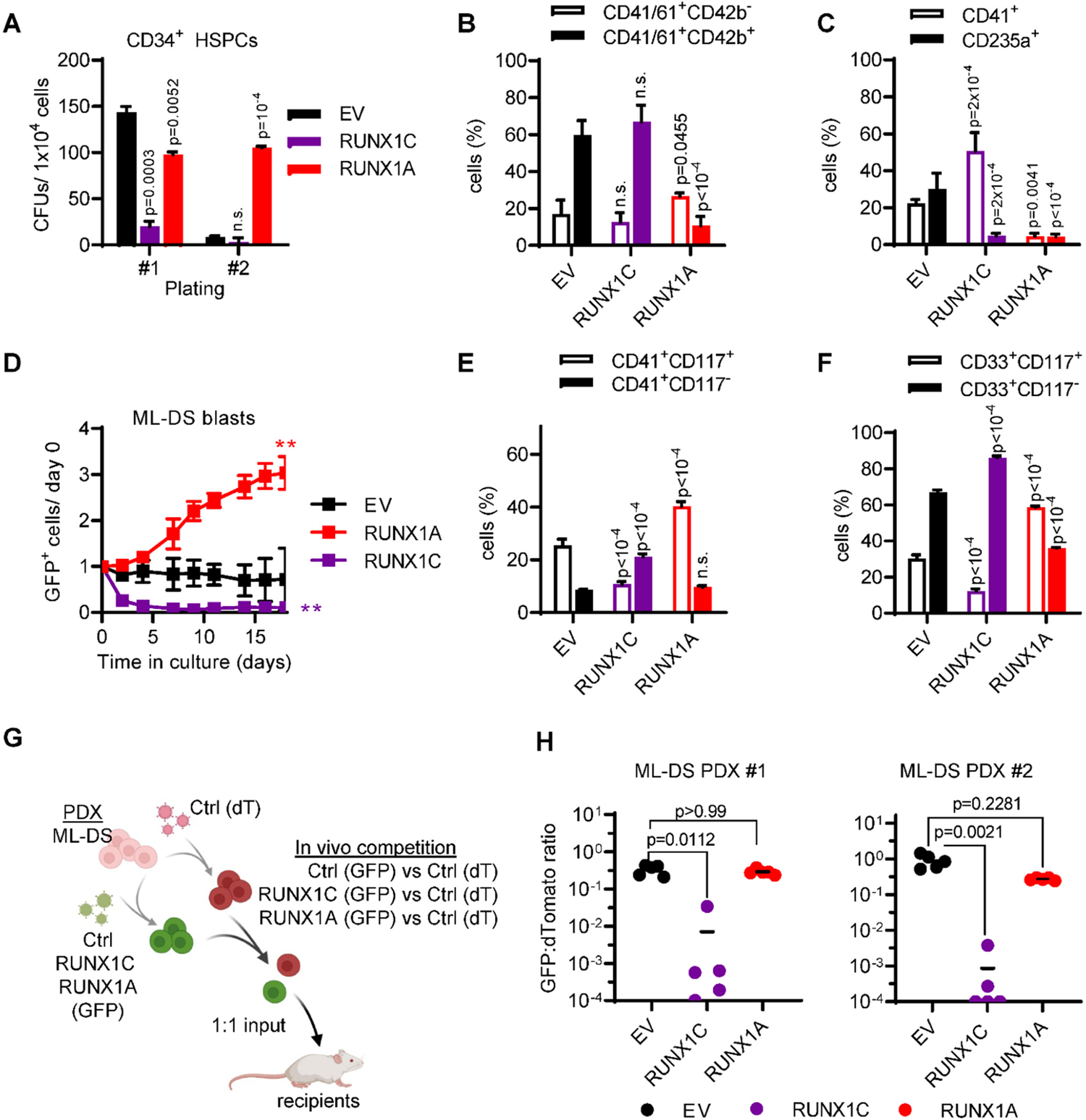
Increased RUNX1A:RUNX1C ratio induces malignant ML-DS phenotype. (A) Colonies after plating *RUNX1A*-, *RUNX1C*- or empty vector control (EV)-transduced human neonatal CD34^+^ HSPCs from two independent donors in methylcellulose-based CFU assays (mean±s.d., n=2; one-way ANOVA). (B-C) Human CD34^+^ HSPCs were lentivirally transduced with *RUNX1A*, *RUNX1C* or empty vector control (EV) (mean±s.d., n=5, two-way ANOVA). (B) Percentage of mature megakaryocytes (CD41^+^CD61^+^/CD42^+^) after 7 days in media promoting megakaryocytic differentiation. (C) Percentage of megakaryocytic (CD41^+^) and erythroid cells (CD235a^+^) after 7 days in media promoting erythroid and megakaryocytic differentiation. (D-F) ML-DS patient derived cells were lentivirally transduced with *RUNX1A*, *RUNX1C* or empty vector control (EV). (D) Percentage of GFP^+^ transduced cells normalized to day 0 post transduction (mean±s.d., n=3, one-way ANOVA, **P_ANOVA_<0.01). Bar graphs showing the percentage of (E) immature CD41^+^CD117^+^ and mature CD41^+^CD117^-^ megakaryocytic cells or (F) immature CD33^+^CD117^+^ and mature CD33^+^CD117^-^ myeloid cells 5 dayspost transduction (mean±s.d., n=3, two way ANOVA). (G) Experimental setup for evaluating RUNX1A:RUNX1C restoration *in vivo*. ML-DS blasts from two patients were transduced with *RUNX1A* (GFP^+^), *RUNX1C* (GFP^+^) or empty vector (EV) control (GFP^+^) and mixed 1:1 with EV control-transduced blasts (dTomato^+^), before transplantation into sub-lethally irradiated recipient mice. (H) Ratio of GFP^+^ to dTomato^+^ cells in the bone marrow (BM) of mice sacrificed 4-8 month after transplantation (n=5, Kruskal-Wallis test). See also **Figure S2** and **Table S2**.

Importantly, restoring the *RUNX1A*:*RUNX1C* equilibrium through *RUNX1C* expression in ML-DS patient-derived blasts (**Table S2**) halted proliferation and accelerated differentiation, as indicated by loss of CD117 and gain of CD33 and CD41 expression (**Figure 2D-F** and **Figure S2F-H**). Inversely, ectopic expression of *RUNX1A* conferred a growth advantage in ML-DS blasts, and led to an accumulation of cells with an immature phenotype (CD33^+^CD117^+^ or CD41^+^CD42b^-^CD117^+^ myeloid/megakaryocytic blasts; **Figure 2D-F** and **Figure S2F-H**). Lastly, we evaluated restoration of *RUNX1C* expression *in vivo* through fluorescence-based competitive transplantation assays using two ML-DS patient-derived xenografts (PDX; **Figure 2G** and **Table S2**). In both cases, *RUNX1C*-expressing leukemic blasts were significantly diminished in the bone marrow of recipient mice at the experimental endpoint, whereas *RUNX1A*-expressing leukemic blasts were unchanged compared to control-transduced blasts (**Figure 2H**).

In summary, these experiments in primary human cells depict a landscape of perturbed differentiation and accelerated proliferation guided by *RUNX1A*. Our data further demonstrate that restoring *RUNX1A:RUNX1C* equilibrium can overcome differentiation arrest in ML-DS blasts, leading to their depletion.

### *RUNX1A* synergizes with *Gata1s* in the leukemic transformation of fetal liver cells

As TAM occurs *in utero*, implying a fetal origin for this disease, we employed a murine fetal liver cell (FLC)-based *in vitro* model to study the oncogenic potential of *RUNX1* isoform imbalance in concert with mutated *Gata1s* (Labuhn *et al*., 2019). Notably, mice do not express the *RUNX1A* isoform (Komeno *et al*., 2014), underlining important species-specific differences that could explain the inability of previous Down syndrome mouse models to recapitulate the human TAM/ML-DS phenotype (Klusmann *et al*., 2010; Li et al., 2005). Combined lentiviral transduction of Cas9-knock-in FLCs with *RUNX1A* and a sgRNA targeting exon 2 of *Gata1* – thereby introducing *Gata1s* mutations – indeed resulted in a robust hyperproliferative phenotype (4140-fold; **Figure 3A**) compared to cells transduced with a combination of a non-targeting control sgRNA (sgCtrl) and empty vector (EV). Individually, *Gata1s* (sgGata1+EV) and *RUNX1A* expression (sgCtrl+*RUNX1A*) also enhanced growth (**Figure 3A**; 366-fold and 140-fold, respectively), but the effect was less pronounced than in combination, suggesting synergy between *Gata1s* and *RUNX1A*. Inversely, ectopic expression of *RUNX1C* alone or in combination with *Gata1s* mutations resulted in a growth disadvantage (**Figure 3A**). We note that none of the other Hsa21 candidate genes from our sgRNA dropout screen produced a synergistic hyperproliferative phenotype with *Gata1s* (**Figure S3A**). Immunophenotypically, the hyperproliferative *RUNX1A Gata1s*-FLCs were CD117^+^CD41a^+^ and partially CD71^+^Ter119^+^ (**Figure 3B**), corresponding to megakaryocytic progenitors with erythroid features – a hallmark of TAM/ML-DS. The expression of erythroid markers was increased upon *RUNX1A* expression compared to control *Gata1s*-FLCs (**Figure 3B**). The synergistic effect of *RUNX1A* and *Gata1s* on the proliferation of FLCs was also observed in culture conditions promoting the expansion and differentiation of erythroid progenitor cells, but not in conditions promoting myeloid expansion and differentiation (**Figure S3B-C**).

**Figure 3.**
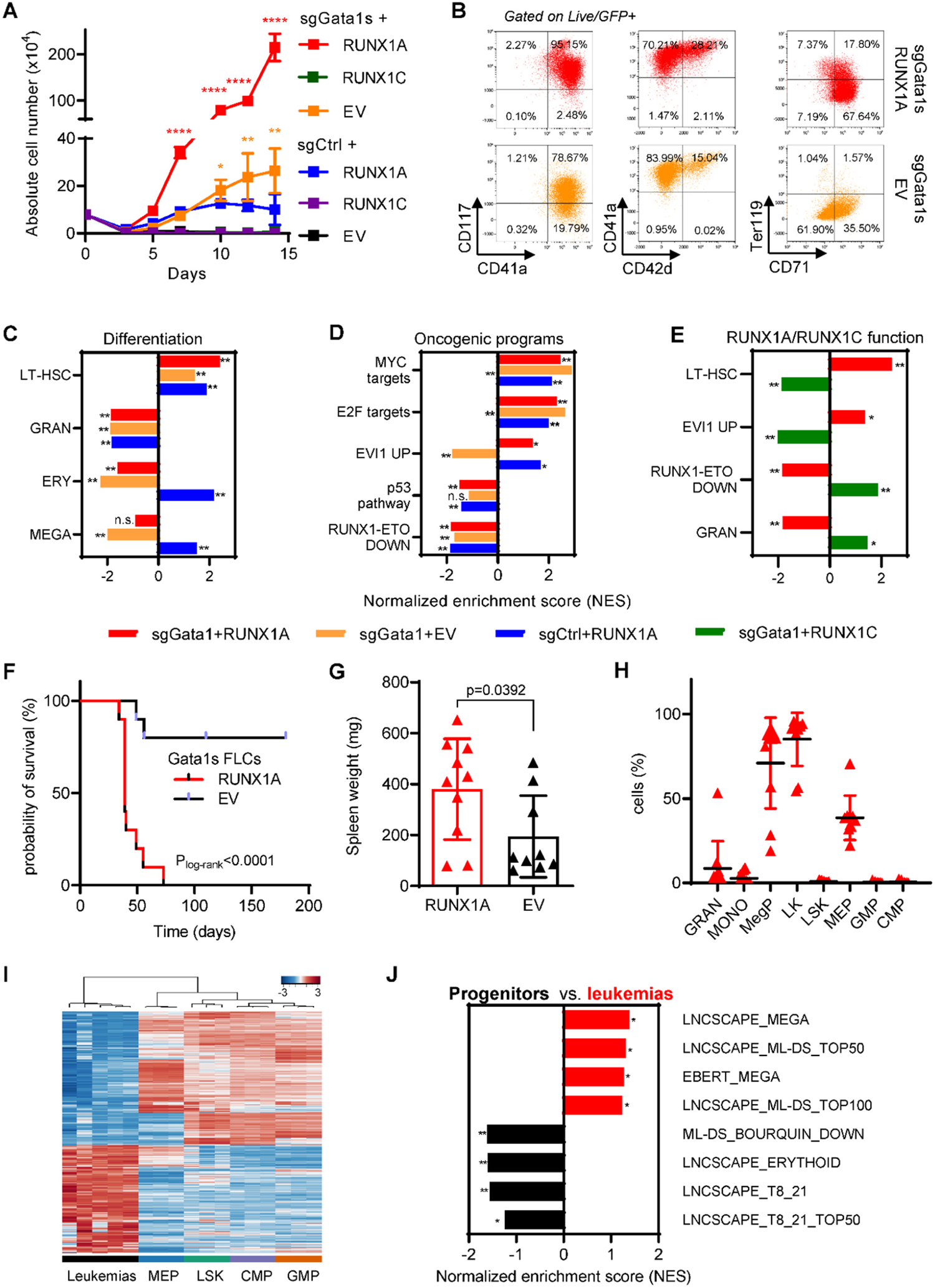
*RUNX1A* synergizes with *Gata1s* in leukemic transformation of murine fetal liver cells. Cas9-knock-in Ter119^-^ FLCs were lentivirally transduced with either *Gata1* (sgGata1s) or control (sgCtrl) sgRNAs, as well as with *RUNX1A, RUNX1C* or empty vector control (EV). (A) Absolute cell number of murine FLCs after combined transduction with lentiviral vectors and maintenance in liquid cultures. Data from one representative experiment performed in replicates are shown as mean±s.d (two-way ANOVA). *P_ANOVA_<0.05, **P_ANOVA_<0.001; ****P_ANOVA_<0.0001. (B) Representative flow cytometry plots of murine sgGata1s+*RUNX1A* or sgGata1s+EV transduced FLCs in liquid culture. The percentage of cells belonging to each immunophenotype is indicated in the corresponding gates. (C-D) Bar graphs showing normalized enrichment scores (NES) of significantly up- or downregulated gene sets associated with differentiation and oncogenic cellular programs, in *Gata1s-* (sgGata1s+EV, orange), *RUNX1A-* (sgCtrl+*RUNX1A*, blue) and *Gata1s-* and *RUNX1A-* (sgGATA1s+*RUNX1A*, red) murine FLCs compared to sgCtrl+EV control FLCs. This analysis highlights the unique and synergistic functional features of *Gata1s* and *RUNX1A*. *FDR q-value<0.25; **FDR q-value<0.05. (E) Bar graph showing normalized enrichment scores (NES) of significantly up- or downregulated gene sets in *Gata1s*+*RUNX1A* (red) and *Gata1s*+*RUNX1C* (green) expressing FLCs compared to control sgCtrl+EV FLCs. This analysis highlights the contrasting functional features of *RUNX1A* and *RUNX1C* on a *Gata1s* background. *FDR q-value<0.25; **FDR q-value<0.05. (F-H) Analysis of mice transplanted with *RUNX1A*- or EV-transduced *Gata1s*-FLCs (n=10 per group), including comparisons of (F) Kaplan-Meier survival curves (log-rank test), (G) spleen weights (unpaired t-test), and (H) flow cytometry on bone marrow cells from the diseased mice (two-way ANOVA). (I) Heatmap representation of RNA-seq data, showing unsupervised hierarchical clustering on the 2,824 most variable genes (standard deviation >1) across five bone marrow samples from leukemic mice and FACS-sorted murine fetal liver HSCs [LSK (Lin^-^Sca1^+^cKit^+^), common myeloid progenitors (CMP) (Lin^-^Sca1^-^cKit^+^CD34^+^ CD16/32^med^), granulocyte-monocyte progenitors (GMP) (Lin^-^Sca1^-^cKit^+^CD34^+^ CD16/32^+^), and megakaryocyte-erythroid progenitors (MEP) (Lin^-^Sca1^-^cKit^+^CD34^-^ CD16/32^low^)]. The sample types are color-coded on the bottom of the heatmap and Z-scores are indicated by the legend at the top of the panel. (J) Bar graph showing normalized enrichment scores (NES) of significantly up- or downregulated gene sets in the murine leukemia samples compared to all progenitor populations. *FDR q-value<0.25; **FDR q-value<0.05. See also **Figure S3** and **Table S3-4**.

Comparative transcriptomic analysis of RNA-sequencing (RNA-seq) data substantiated the synergy between *Gata1s* mutations and *RUNX1A* expression. As previously described, gene set enrichment analysis (GSEA) revealed a marked reduction of erythroid genes and concurrent activation of pro-proliferative genes – including MYC and E2F targets – in *Gata1s* mutated FLCs (**Figure 3C-D** and **Table S3**) (Alejo-Valle *et al*., 2021; Klusmann *et al*., 2010; Ling et al., 2019). While *RUNX1A* expression mitigated the reduction of erythroid genes caused by *Gata1s*, it triggered a similar upregulation of MYC and E2F target genes, suggesting convergence on these oncogenic pathways. In addition, *RUNX1A* expression induced target genes of EVI1 – one of the most invasive proto-oncogenes in human leukemia – and a long-term hematopoietic stem cell signature, whereas the opposite was true upon *RUNX1C* expression (**Figure 3D-E** and **Table S3**). Hence, the synergistic oncogenic expression program of *Gata1s* and *RUNX1A* is characterized by the induction of EVI1, MYC and E2F genes target genes, and a long-term hematopoietic stem cell signature, and the concomitant repression of erythroid and megakaryocytic differentiation signatures.

To further explore the leukemogenic potential of *RUNX1* isoform disequilibrium in the pathogenesis of Down syndrome-associated leukemia, we performed *in vivo* experiments using *Gata1s*-FLCs. Upon transplantation into sub-lethally irradiated syngeneic recipients (C57Bl/6J), *Gata1s*-FLCs typically become transiently abundant in the peripheral blood (Labuhn *et al*., 2019). In contrast, *RUNX1A*-expressing *Gata1s*-FLCs caused high penetrance (100%) leukemia with a short latency (median survival: 39 days) and organ infiltration (**Figure 3F-G** and **Figure S3D**). Detailed flow cytometry analysis revealed cells of a megakaryocytic progenitor-like phenotype (CD41^+^CD117^+^CD34^-^CD16/32^low^) resembling TAM and ML-DS (**Figure 3H** and **Figure S3E**), which re-initiated disease when transplanted into secondary recipients (**Figure S3F**). Notably, *RUNX1A*-expression did not cause leukemia in wild-type *Gata1*-FLCs (**Figure S3G**). Transcriptomic analysis of RNA-seq data from the murine leukemias in comparison to stringently sorted murine fetal liver stem and/or progenitor populations confirmed the megakaryocytic progenitor-like phenotype and the ML-DS-like gene expression profile (Bourquin *et al*., 2006; Schwarzer et al., 2017) (**Figure 3I-J** and **Table S4**). Importantly, Cre recombinase-mediated excision of the LoxP-flanked *RUNX1A* cDNA induced rapid depletion of the leukemic blasts *in vitro*, underlining their dependency on *RUNX1A* and its importance in ML-DS pathogenesis (**Figure S3H**). Of note, miR-125 further accelerated the leukemic transformation of *Gata1s*-FLCs by *RUNX1A,* suggesting synergy between RUNX1A and another well characterized oncogene on chromosome 21 (Alejo-Valle *et al*., 2021; Emmrich et al., 2014) (**Figure S3I**).

These data demonstrate that the interplay between *Gata1s* and *RUNX1* isoform disequilibrium results in the proliferation and accumulation of immature megakaryocytic progenitors *in vitro*, and in ML-DS-like leukemia or an aggressive form of TAM *in vivo*.

### Distinct RUNX1A and RUNX1C protein interaction networks

To better understand the oncogenic mechanisms mediated by RUNX1A in TAM/ML-DS, we compared the RUNX1A and RUNX1C protein interaction networks via co-immunoprecipitation of doxycycline-inducible HA-tagged RUNX1A and RUNX1C in CMK cells, followed by mass spectrometry of the bound cofactors (**Figure 4A**). 98 and 57 proteins were significantly bound by RUNX1C and RUNX1A, respectively (**Figure 4B** and **Figure S4A**). Of these, 45 proteins were commonly bound by both RUNX1 isoforms, including CBFβ and other *bona fide* RUNX1 interaction partners (**Figure 4B** and **Figure S4A**). 53 protein interactions were unique to RUNX1C. RUNX1C-specific cofactors were enriched for proteins involved in the regulation of cell cycle, gene expression, protein and mRNA metabolism as well as chromosome organization. Interestingly, members of the spliceosome A/C and NSL complexes (e.g. SF3B1, WDR5 and KANSL2) were among the RUNX1C-specific cofactors (**Figure S4B**), suggesting an active role in splicing and chromosomal organization that is lost in RUNX1A, as demonstrated for the RUNX1::RUNX1T1 fusion oncoprotein (Grinev et al., 2021). In contrast, the 12 RUNX1A-specific cofactors are involved in active transcription/replication and G1-S transition (**Figure 4B**). Importantly, MAX – a crucial cofactor of MYC – was among the RUNX1A-specific interacting proteins, as verified via Western blot (**Figure 4C**). GATA1s co-immunoprecipitated with RUNX1A and RUNX1C, albeit not at a significant enrichment level (data not shown). As GATA1 is a well-described RUNX1 interaction partner (Elagib et al., 2003) and as *GATA1s* mutations characterize TAM/ML-DS (Alford et al., 2011; Kanezaki et al., 2010), we investigated the putative differential binding of RUNX1A and RUNX1C in more detail. To this end, we pulled down doxycycline-induced HA-tagged GATA1 or GATA1s in stably transduced CMK cells, which harbor endogenous *GATA1s* mutations and hence exclusively express GATA1s. This is an important point, since GATA1 forms homodimers (Crossley et al., 1995; Shimizu et al., 2007) precluding the enrichment of GATA1s-specific protein complexes in the presence of GATA1. We found that both GATA1 and GATA1s interact with RUNX1C, as previously described (Xu et al., 2006). However, neither GATA1 nor GATA1s appear to interact with RUNX1A (**Figure 4D**). Thus, our proteomic data suggest an altered protein interaction network, which may contribute to the TAM/ML-DS specific phenotype of RUNX1A in *GATA1*-mutated cells.

**Figure 4.**
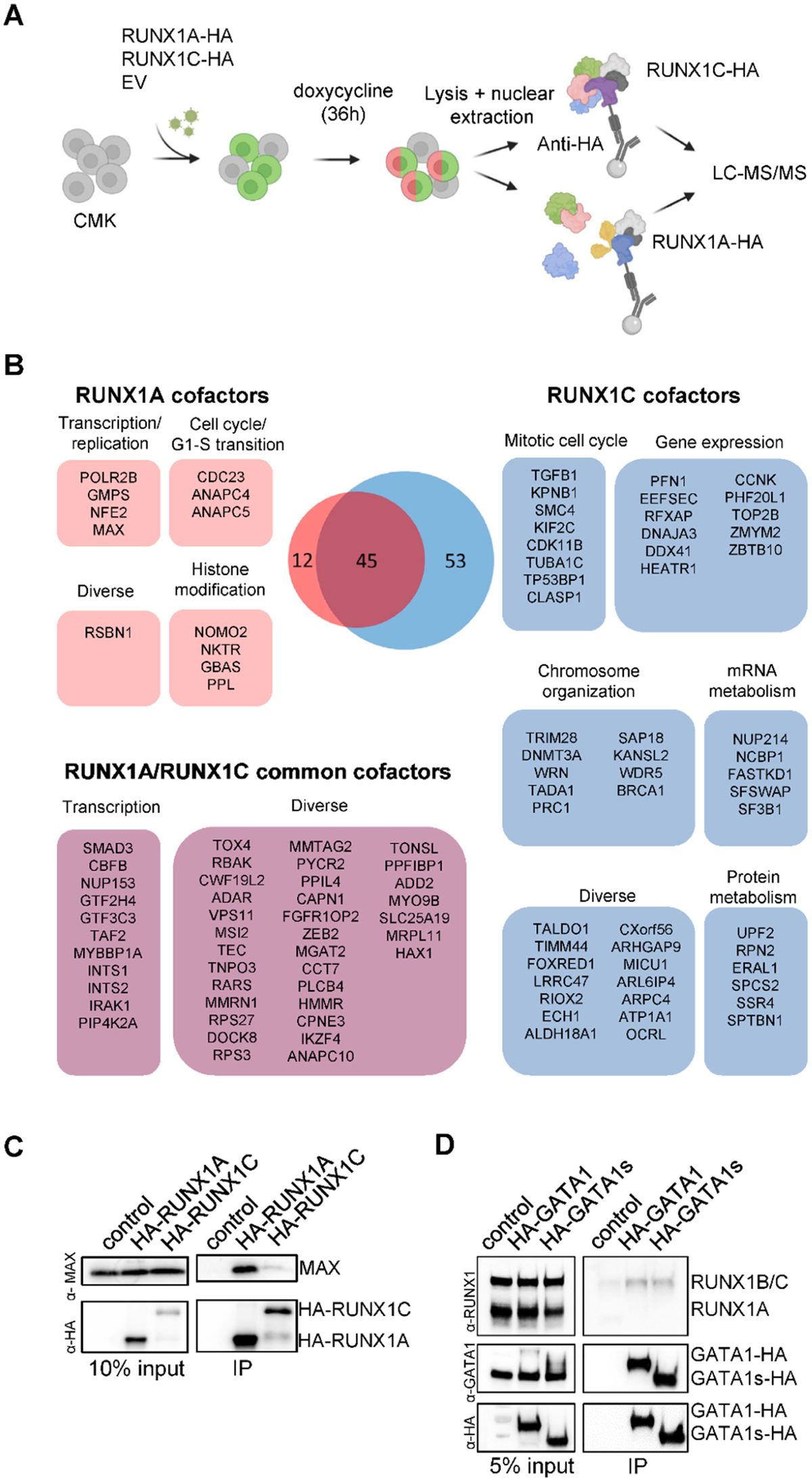
Distinct RUNX1A or RUNX1C protein interaction networks. (A) Experimental setup for isolating HA-RUNX1A- or HA-RUNX1C-containing protein complexes from CMK cells. (B) RUNX1A and RUNX1C complex composition in CMK cells with functional grouping. The Venn diagram shows significantly enriched interactors (log_2_ fold change >1; p-value <0.05). (C) Western blot confirming co-immunoprecipitation of MAX and RUNX1A using anti-MAX and anti-RUNX1 antibodies. Representative picture of 2 independent experiments using K562 cells are shown. 10% input was used as the loading control. (D) Western blot showing RUNX1A/C isoforms co-immunoprecipitated with doxycycline-inducible HA-tagged GATA1 and GATA1s or the empty control vector. Representative picture of 3 independent experiments using CMK cells are shown. 5% input was used as the loading control. See also **Figure S4.**

### RUNX1A affects gene regulation by displacing endogenous RUNX1C

To further interrogate the RUNX1A-centered protein interaction network and determine the consequences of its inability to form a complex with GATA1 or GATA1s, we performed CUT&RUN on endogenous GATA1s and RUNX1C (using a C-terminal antibody that does not recognize exogenous RUNX1A) in *Gata1s*-FLC with or without exogenous expression of doxycycline-inducible HA-tagged RUNX1A or RUNX1C. We found that 33% of the promoter or enhancer regions occupied by endogenous RUNX1C were also occupied by HA-RUNX1A (**Figure 5A**), and that half (52%) of the RUNX1C/RUNX1A occupied regulatory regions were co-bound by GATA1s (**Figure 5A**), corroborating the known GATA1-RUNX1 interplay in transcriptional regulation (Tijssen et al., 2011).

**Figure 5.**
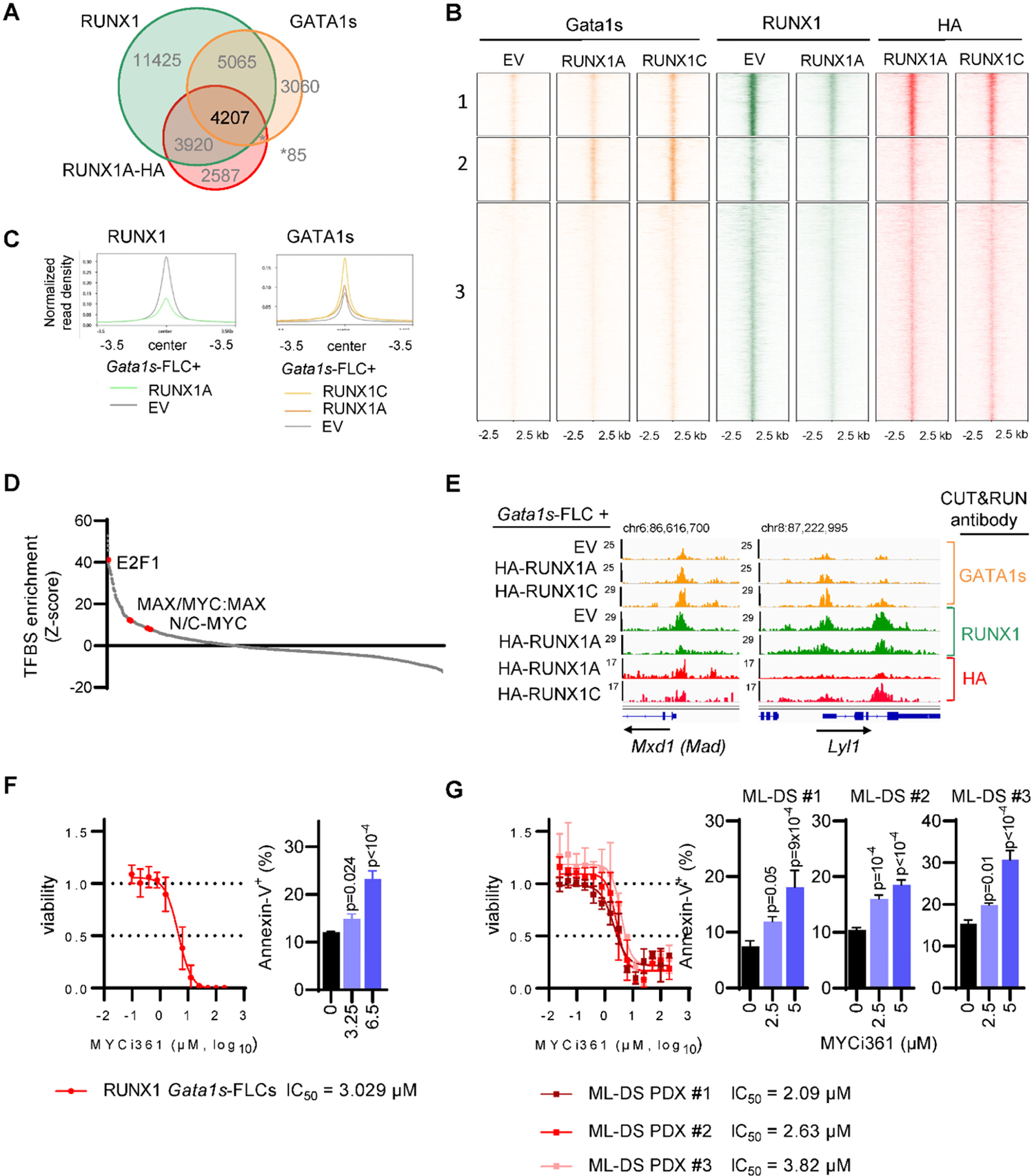
RUNX1A affects gene regulation by displacing endogenous RUNX1C. (A-E) CUT&RUN results from murine *Gata1s*-FLCs after doxycycline-induced overexpression of HA-RUNX1A and HA-RUNX1C as well as EV control. (A) Venn diagram showing the number of genomic regions bound by endogenous RUNX1, endogenous GATA1s and/or HA-RUNX1A after HA-RUNX1A overexpression. Peaks were called by SEACR (Meers et al., 2019b) using EV control cells as background. (B) Heatmaps depicting the co-localization of endogenous GATA1s (orange, left), endogenous RUNX1 (green, middle), and HA-RUNX1A/-RUNX1C (red, right) signals after doxycycline-induced HA-RUNX1A or HA-RUNX1C expression in Gata1s-FLCs. Empty vector (EV) transduced cells were used as a control. The data were generated via CUT&RUN using the antibodies indicated above the graphs. Regions ±2.5 kb of the peak center are shown. (C) Binding intensities (normalized read density) of RUNX1 (left) and GATA1s (right), in EV control and doxycycline-induced HA-RUNX1A and HA-RUNX1C expressing cells. (D) Transcription factor motif enrichment analysis under RUNX1 peaks, at the promoter regions of genes that are differentially expressed upon doxycycline induced RUNX1A expression (log_2_ FC = 1). Human promoter regions were used as background. Z-score intensities are shown. (E) IGV snapshots of *Mxd1* and *Lyl1* gene promoters showing occupancy of endogenous GATA1s, RUNX1 and HA-RUNX1A/-RUNX1C in *Gata1s*-FLCs after doxycycline induced HA-RUNX1A/-RUNX1C expression or in EV control expressing cells. The tracks display coverage (RPKM) (left). Scale and chromosome location are shown (top). (F) Dose response curves for MYC:MAX dimerization inhibitor MYCi361 on RUNX1A *Gata1s*-FLCs 24h after treatment *in vitro* (left). IC_50_-values are depicted below the graph. Bar graph showing the percentage of AnnexinV^+^ cells after treatment with the indicated doses of MYCi361 in comparison to the DMSO control (right; mean ± SD; n=3; one-way ANOVA). (G) Dose response curves for MYC:MAX dimerization inhibitor MYCi361 on ML-DS blasts derived from three patients 24h after treatment *in vitro* (left). IC_50_-values are depicted below the graph. Bar graph showing the percentage of AnnexinV^+^ cells after treatment with the indicated doses of MYCi361 in comparison to the DMSO control (right; mean ± SD; n=3; one-way ANOVA). See also **Figure S4-5** and **Table S5**.

Unsupervised clustering of peaks at regulatory regions bound by endogenous RUNX1C, GATA1 and HA-RUNX1A revealed three clusters (**Figure 5B**). *De novo* motif discovery uncovered an enrichment of RUNX family motifs in clusters 1 and 3, whereas GATA family motifs were most abundant in cluster 2 (**Figure S4C**). Importantly, we observed a global reduction of endogenous RUNX1 occupancy upon HA-RUNX1A expression across all clusters (**Figure 5C** and **Figure S4D**). Consistent with our protein-protein interaction studies, GATA1s binding increased globally upon expression of HA-RUNX1C, in comparison to HA-RUNX1A and the empty vector control (**Figure 5C** and **Figure S4E)**.

Interestingly, the majority of the genes in the three CUT&RUN clusters were upregulated upon HA-RUNX1A expression (67.6-81.5%) (**Figure S4F**). Transcription factor motif analysis of peaks in the promoter regions of differentially expressed genes revealed an overrepresentation of the MYC-coactivation factor E2F as well as of single MYC, MAX and MYC:MAX dimeric motifs (**Figure 5D** and **Table S5**). Among the RUNX1A-induced genes with MAX or MYC:MAX binding motifs in their promoter regions are *Mxd1* (*Mad1; Max Dimerization Protein 1*), *Myct1 (MYC target 1)*, known oncogenic drivers such as *Lyl1* and *Malat1* as well as the megakaryocyte gene *Itga2b* (CD41) – all of which were repressed by RUNX1C in RNA-seq experiments (**Figure 5E** and **Figure S4G-I)**. These data are consistent with our comparative transcriptomic analyses in *Gata1s*-FLCs, which uncovered the activation of MYC and E2F targets by GATA1s and RUNX1A resulting in a synergistic induction of proliferation and an immature phenotype, while the opposite was true for *RUNX1C* (**Figure 3C-E**).

Taken together, our proteomic and genomic analyses imply that RUNX1A interferes with normal RUNX1C function by displacing it from target gene promoters. Instead, RUNX1A interacts with the MYC co-factor MAX and induces an MYC- and E2F-driven expression program. The concerted alteration of normal megakaryocytic progenitor gene expression programs through GATA1s and RUNX1A leads to malignant transformation in these cells, via the upregulation of oncogenic gene expression programs and the perturbation of RUNX1C-regulated differentiation programs, respectively.

### Targeting MYC:MAX dimerization as a therapeutic approach for ML-DS

Finally, we investigated the role of the MYC co-factor MAX, which we hypothesized to be central to the synergy between GATA1s and RUNX1A, during leukemia pathogenesis. ShRNA-mediated knockdown and sgRNA-mediated knockout of *MAX* inhibited the growth of CMK cells *in vitro* (**Figure S5A-B**). These findings were confirmed in murine *RUNX1A Gata1s*-FLCs and in ML-DS blasts derived from two patients (**Figure S5C-F** and **Table S2**). Normal CD34^+^ HSPCs showed only a mild growth impairment upon *MAX* depletion (**Figure S5G-H**). Importantly, pharmacological disruption of MYC:MAX dimerization using the MYC inhibitor MYCi361 (Han et al., 2019) caused apoptotic cell death in *RUNX1A Gata1s*-FLCs in a dose dependent manner (**Figure 5F**). Accordingly, MYCi361 also induced apoptosis in ML-DS blasts derived from three patients, with median lethal concentration (IC_50_) values of 2.09 µM, 2.63 µM and 3.82 µM, respectively (**Figure 5G**). Normal CD34^+^ HSPCs from two donors had IC_50_ values of 7.291 µM and 7.168 µM, respectively (**Figure S5I**), outlining a therapeutic window. Thus, we demonstrate that the MYC co-factor MAX is key to leukemic transformation mediated by RUNX1A and GATA1s, and can be therapeutically exploited.

## Discussion

Aneuploidies, and in particular partial or complete amplifications of chromosome 21, are frequent alterations in leukemia; however, their underlying pathomechanisms remain enigmatic (Ben-David and Amon, 2020). In this study, we leveraged the model system of Down syndrome-associated TAM and ML-DS to interrogate oncogenic factors on chromosome 21 for roles in AML pathogenesis. A CRISPR-Cas9 screen of chromosome 21 genes unexpectedly revealed *RUNX1* as an ML-DS dependency in cell lines and patient-derived blasts *in vitro* and *in vivo*, despite former controversy regarding its role in TAM and ML-DS (Kirsammer *et al*., 2008; Maclean et al., 2012). Through detailed functional validation, we discovered that, rather than *RUNX1* gene dosage, a disequilibrium of *RUNX1* isoforms and *RUNX1A* bias is key to trisomy 21-associated leukemogenesis. By dissecting the consequences of *RUNX1* disequilibrium in TAM/ML-DS, we showed that *RUNX1A* acts in concert with pathognomonic *GATA1s* mutations in this context, thereby blocking megakaryocytic differentiation and accelerating progenitor proliferation – effects that were reversed upon restoring *RUNX1A*:*RUNX1C* equilibrium in murine models and patient-derived blasts *in vitro* and *in vivo*. These findings are in line with the known function of *RUNX1C* during adult hematopoiesis (Growney et al., 2005; Ichikawa et al., 2004; Senserrich et al., 2018), where it regulates megakaryopoiesis by controlling the proliferation and survival of committed progenitors (Draper et al., 2017). Thus, through the systematic interrogation of chromosome 21-encoded coding genes, our work contributes to understanding the synergy between trisomy 21 and *GATA1* mutations in ML-DS leukemogenesis, wherein RUNX1A can further enhance the oncogenic effect of miR-125b, another well-characterized oncogenic driver on chromosome 21 (Alejo-Valle *et al*., 2021). We suggest a model where *RUNX1A*:*RUNX1C* disequilibrium in TAM and ML-DS is a consequence of trisomy 21 in fetal HSPCs and the impaired capacity of GATA1s to repress *RUNX1A*. Our findings have further implications for other types of leukemia with numerical or structural alterations of chromosome 21 and/or *RUNX1* isoform disequilibrium, the latter of which we observed in non-DS acute megakaryoblastic leukemia patient-derived xenografts (**Figure S1D-F**). However, different mechanisms may contribute to *RUNX1A:RUNX1C* dysregulation in other subtypes of AML or myelodysplastic syndrome (MDS) (Sakurai et al., 2017).

Mechanistically, we showed that the altered protein interaction network of RUNX1A (compared to RUNX1B/C) underlies its oncogenic function. RUNX1A fails to interact with GATA1 and GATA1s to control megakaryopoiesis (Xu *et al*., 2006). Rather, our proteomics analyses point towards RUNX1A’s increased affinity for the MYC cofactor MAX. We propose a model where RUNX1A, in complex with MAX, displaces RUNX1C from its endogenous binding sites, where it normally recruits GATA1. Thereby, RUNX1A perturbs RUNX1C:GATA1 regulated gene expression and activates MYC:MAX and E2F target genes, leading to a megakaryocytic differentiation block and acceleration of progenitor proliferation. This model would help explain the synergistic oncogenic effect of RUNX1A and GATA1s. GATA1s not only fails to repress *RUNX1A* but E2F and MYC target genes during megakaryopoiesis (Klusmann *et al*., 2010), suggesting a vicious circle in fetal trisomy 21 HSPCs. The central role of MAX in the RUNX1A-induced phenotype was validated by genetically interfering with *MAX* expression. Importantly, the dependency of ML-DS blasts on intact MAX represents a vulnerability that can be therapeutically exploited, as we demonstrated by using the MYC:MAX dimerization inhibitor MYCi361 (Han *et al*., 2019) to eradicate ML-DS patient blasts.

Although previous studies have reported the oncogenic and dominant negative effects RUNX1A exerts on RUNX1B (Bertrand-Philippe et al., 2004; Guo et al., 2012; Ran et al., 2013; Sakurai *et al*., 2017; Tanaka et al., 1995), our findings are unexpected and have general implications for basic oncology research. First, our study demonstrates that species-specific differences in isoform expression must be considered when interpreting species-specific or divergent phenotypes in mouse models. *RUNX1A* is a primate-specific isoform of *RUNX1* (Komeno *et al*., 2014), which may explain the inability of Down syndrome mouse models (Alford et al., 2010; Li *et al*., 2005) to recapitulate the human TAM/ML-DS phenotype or the interplay of GATA1s and trisomy 21 seen in human induced pluripotent stem cells (Byrska-Bishop et al., 2015; Chou *et al*., 2012; Maclean *et al*., 2012; Roy *et al*., 2012). Second, our study underlines the importance of accounting for all isoforms when studying the oncogenic effects of a given gene and emphasizes the importance of alternative splicing in the pathogenesis of cancer. Various mutations in splicing factors have been identified in AML and MDS genomes (Makishima et al., 2012). Whether these mutations are causative for the increased RUNX1A:RUNX1B/C ratio observed in MDS (Sakurai *et al*., 2017) will need to be investigated in the future.

Overall, our study provides a general framework for interrogating the contribution of aneuploidy to oncogenesis. Given the current state of the field, in which we have a near-complete map of the cytogenetic, mutational and epigenetic landscape of cancer, these insights will be crucial for understanding the complex cooperation between common fusion oncogenes, recurrently mutated genes and larger amplifications or deletions during disease initiation and progression. As we illustrate with the identification of MAX as a RUNX1A co-factor and its pharmacological inhibition, this knowledge can have direct therapeutic implications.

## Supporting information

Key Resource Table

## Acknowledgements

We thank D. Trono of EFPL, Lausanne, Switzerland, for kindly providing pMD2.G (Addgene plasmid 12259) and psPAX2 (Addgene plasmid 12260) and D.E. Zhang for sharing the *RUNX1A* cDNA. J.H.K receives funding from the European Research Council (ERC) under the European Union’s Horizon 2020 research and innovation programme (grant agreement No 714226) and is a recipient of the St. Baldrick’s Robert Arceci Innovations Award. D.H was supported by the Deutsche Krebshilfe (DKH; #111743). This work was supported by grants to J.H.K. from the German Research Foundation (DFG; KL-2374/1-3) and the DKH (#109251 and #110806). S.G., M.L., F.J.S., S.K.K. and L.S. were supported by the Hannover Biomedical Research School.

## Author contributions

S.G. and D.B.H. performed experiments, analyzed and interpreted data and revised the manuscript. R.B., O.A.V., H.I., C.B., M.L., C.I., L.S, F.J.S., S.K.K. and. R.T.V. performed experiments and analyzed data. E.R., M.G, S.M, V.A and T.R. performed bioinformatics analysis and data interpretation, and revised the manuscript. A.S., S.H., M.L.Y. supervised the analyses and revised the manuscript. D.R. provided patient material and data. D.H. and J.H.K. designed the study, analyzed and interpreted data, wrote the manuscript and academically drove the project.

## Methods

### Resource availability

#### Lead contact

Further information and requests for resources and reagents should be directed to and will be fulfilled by the lead contact, Jan-Henning Klusmann.

#### Materials availability

Plasmids generated in this study have been deposited to Addgene (see Key Resources Table).

#### Data availability

RNA-Seq gene expression data have been deposited in the European Nucleotide Archive (ENA) and are accessible through accession numbers: ERS6071576 - ERS6071595. CUT&RUN data are accessible through ENA with the accession numbers: ERS5956460 - ERS5956467. Mass spectrometry proteomics data have been deposited to the ProteomeXchange Consortium via the PRIDE partner repository with the dataset identifier identifier PXD030616 and 10.6019/PXD030616. Data will be available immediately following publication, no end date. Other remaining data are available within the Article and Supplementary Files, or from the authors upon request.

### Experimental model and subject details

#### Cells and cell culture

All human cell lines used for the purposes of this publication (CMK, K562 and HEK293T) were obtained from the German Collection of Microorganisms and Cell Cultures (DSMZ, Braunschweig). Culturing and maintenance were performed according to the supplier’s instructions. Murine fetal liver cells were isolated from E12.5-E13.5 *Cas9* heterozygous C57BL/6J (Jackson Laboratory) embryos upon depletion of erythroid cells using anti-Ter119 immunomagnetic microbeads (StemCell Technologies). Neonatal CD34^+^ HSPCs were obtained from cord blood of healthy donors and enriched using anti-CD34 immunomagnetic microbeads (Miltenyi Biotech and StemCell Technologies). All investigations were performed in accordance to the Declaration of Helsinki and informed consent was obtained according to local laws and regulations. Murine FLCs were stimulated before transduction in DMEM (Gibco, Life Technologies) with 10% FCS (Capricorn Scientific), 1% streptomycin/penicillin (Millipore), 1% L-glutamine (Millipore), 1% sodium pyruvate (Gibco, Life Technologies), 1% non-essential amino acids (Gibco, Life Technologies), 25ng/mL Scf and 25ng/mL Tpo, and were further expanded in the same medium except with 2ng/mL Scf. The same medium with low Scf conditions was used in megakaryocytic differentiation assays. For erythroid differentiation, the same cells were expanded in IMDM (Lonza Bioscience) with 15% FCS, 2% streptomycin/penicillin, 1% L-glutamine, 2U/mL Epo, 10μM dexamethasone (Sigma-Aldrich), 5ng/mL Scf and 1ng/mL Il3. For myeloid differentiation, IMDM with 10% FCS, 2% streptomycin/penicillin, 1% sodium pyruvate, 20ng/mL Scf, 40ng/mL G-CSF, 40ng/mL M-CSF, 10ng/mL Il3 and 10 ng/mL Il6 was used. Human HSPCs were expanded in StemSpan SFEM (StemCell Technologies) with 1% streptomycin/penicillin, 50ng/mL SCF, 50ng/mL FLT3-ligand, 20ng/mL TPO, 10ng/mL IL3 and 10ng/mL IL6. For megakaryocytic differentiation, StemSpan SFEM with 1% streptomycin/penicillin, 0.25% CD-liipid concentrate (Gibco, Life Technologies), 30ng/mL TPO, 1ng/mL SCF, 7.5ng/mL IL6 and 13.5ng/mL IL9 was used. For combined megakaryocytic/erythroid differentiation, cells were cultured in StemSpan SFEM with 1% streptomycin/penicillin, 1% L-glutamine, 0.25% CD-lipid concentrate, 50ng/mL SCF, 0.08U/mL EPO, 100ng/mL TPO, 10ng/mL IL3 and 10ng/mL IL6 for 7 days. On day 7, the concentrations of SCF and EPO were increased to 100ng/mL and 0.5U/mL, respectively, while the TPO concentration was reduced to 50ng/mL. For myeloid differentiation, the culture medium consisted of RPMI (Lonza Bioscience), with 10% FCS, 1% streptomycin/penicillin, 1% L-glutamine, 5ng/mL SCF, 5ng/mL GM-CSF, 10ng/mL G-CSF and 5ng/mL IL3. ML-DS blasts from PDX models were cultured *in vitro* in StemSpan SFEM with 1% streptomycin/penicillin, 0.25% CD-lipid concentrate, 50ng/mL SCF, 50ng/mL FLT3, 20ng/mL TPO, 2.5ng/mL IL3, 10ng/mL IL6, 0.75μΜ Stemregenin1 (SR1) (APExBIO) and 35nM UM171 (APExBIO). Leukemic bone marrow cells obtained from primary recipients were expanded *in vitro* in StemSpan SFEM with 1% streptomycin/penicillin, 10ng/mL Scf, 20ng/mL Tpo and 10ng/mL Il3. All human and murine cytokines were purchased from Peprotech, except for EPO, which was purchased from GoldBio. For drug response curves using MYCi361 (Seleckchem), cell viability was assessed using the CellTiter-Glo® Cell Viability Assay (Promega) according to the manufacturer’s instructions.

#### Lentiviral vector construction and transduction

For the CRISPR/Cas9 screen, a library of 1090 sgRNAs targeting 218 genes on Hsa21 was cloned into the SGL40C.EFS.dTomato vector (Addgene, #69147 and #89395) (Reimer et al., 2017). The sgRNAs targeting *RUNX1 (1-5)* were also cloned into the SGL40C.EFS.dTomato vector, while the sgRNA targeting *Gata1 (sgGata1.4)* was cloned into the SGL40C.EFS.dTomato and the pLeGO-iG2 vector (Addgene, #27341) (Weber et al., 2008). A detailed list of all protospacer and shRNA sequences is provided in **Table S1** and the **Key Resources Table**. shRNAs were designed using the miR-N online tool (Adams et al., 2017) and cloned into the SIN40C.SFFV.tBFP2.miR30n backbone. The *RUNX1A* cDNA was cloned into the pLeGO.iG2 vector. Codon-optimized 5’HA-tagged RUNX1A and RUNX1C cDNAs carrying a single HA-tag at the 5’-end were synthesized at Twist Bioscience and introduced into the pRRL.PPT.SFFV.MCS.IRES.eGFP vector and into the doxycycline-inducible SIN40C.TRE.MCS.IRES.idTomato.PGK.sfGFP.P2A.Tet3G vector system. Additionally, Cre recombinase cDNA sequences were cloned into the SIN40C.SFFV.MCS.IRES.dTomato backbone. All sequences were verified by Sanger sequencing.

Lentiviral particles were generated by co-transfection of HEK293T cells with the vector of interest, pMD2.G and psPAX2 (Addgene, #12259 and #12260) using the polyethylenimine (PEI) method (Bhayadia et al., 2018; Emmrich *et al*., 2014). CMK cells, murine FLCs, CD34^+^ HSPCs, PDX-derived ML-DS blasts and leukemic BM cells were lentivirally transduced as previously described (Gialesaki et al., 2018).

#### CRISPR/Cas9 library screening

6×10^6^ CMK and K562 cells were lentivirally transduced with the library of sgRNAs, in order to achieve a 1000-fold coverage of the library. Both cell lines were previously transduced with the pLKO5d.SFFV.Cas9.BlastR vector, to establish stable expression of the Cas9 nuclease. The cells were maintained in culture and samples were harvested for DNA extraction at an early (day 4 upon transduction) and a late time point (after 18 population doublings). DNA was isolated using the QIAmp DNA Blood Mini Kit (Qiagen). Adapters including sample specific barcodes were added by PCR. Libraries were subjected to next generation sequencing (NGS) for sgRNA quantification.

#### Evaluation of RUNX1 isoforms expression by qRT-PCR

Total RNA was extracted from adult CD34^+^ HSPCs, trisomy 21 CD34+ HSPCs, erythroid cells, megakaryocytes, granulocytes, monocytes, and from AMKL, ML-DS, TAM, t(10;11) AML and t(9;11) AML patient samples, using the Quick-RNA Microprep Kit (Zymo Research), and complementary DNA (cDNA) was then synthesized using the High Capacity cDNA Reverse Transcription Kit (Applied Biosystems, Thermo Fisher Scientific). Quantitative real-time PCR (qRT-PCR) was performed in the StepOnePlus™ Real-Time PCR System, using the TaqMan® Fast Advanced Master Mix (both from Applied Biosystems, Thermo Fisher Scientific).

#### Oxford Nanopore sequencing of RUNX1 transcript isoforms

Total RNA from TAM and ML-DS patient-derived samples were isolated using the Quick-RNA Microprep Kit (Zymo Research). The MicroMACS mRNA isolation kit (Miltenyi Biotech) was used for isolation of polyA^+^-enriched RNA. 1 ng polyA^+^ RNA was processed for Oxford Nanopore full length transcript sequencing using the PCR-cDNA Sequencing Kit (SQK-PCS109; Oxford Nanopore Technologies) according to the manufactureŕs protocol. Sequencing was performed on an Oxford Nanopore MinION Mk1B. Base calling was done with guppy v3.0.6. Raw sequencing reads were first processed with filtlong (https://github.com/rrwick/Filtlong), removing reads shorter than 20 nt and associated with mean quality scores below 80. Subsequently, full-length cDNA reads were extracted using pychopper (https://github.com/nanoporetech/pychopper).

For gene expression analysis, full-length reads were mapped to the human genome (UCSC hg38) with minimap2 (Li, 2018) using the long-read spliced alignment preset (-x splice) and without reporting secondary alignments (--secondary=no). Afterwards, sorting and indexing of mapped reads was performed using samtools (Li et al., 2009). Summarization of mapped reads was done with featureCounts (Liao et al., 2014) applying long read counting mode (-L) and using Ensembl GRCh38.89 (Aken et al., 2017) as the annotation base. Differential gene expression was determined using the R package edgeR (Robinson et al., 2010) – utilizing trimmed mean of M-values (TMM; (Robinson and Oshlack, 2010)) for normalization. Normalized gene expression values were expressed as counts per million mapped reads (CPM). For determination of differential transcript usage, full-length reads were mapped to the human transcriptome (Gencode v29; (Frankish et al., 2019)) with minimap2 using the Nanopore preset (-x map-ont) and without reporting secondary alignments (--secondary=no). Transcript expression was assessed with salmon (Patro et al., 2017) in alignment-based mode and activated options for correction of sequence-specific and GC-content bias. The R-package DRIMseq (Nowicka and Robinson, 2016) was used to perform testing for differential transcript

#### In vitro cell growth assays

In all growth assays, the same standard protocol was followed. Every 2-3 days, a sample of cells was taken to measure cell counts or percentage of fluorescent cells on the CytoFLEX platform (Beckman Coulter, BC) or the FACS Canto Flow Cytometer (BD Biosciences). Competition assays were performed by using a starting transfection efficiency of approximately 50%.

#### In vitro differentiation assays

Upon transduction, 5×10^4^ murine FLCs were cultured in the megakaryocytic differentiation-supporting medium, as described above. Similarly, 3×10^4^ cells were cultured in the erythroid medium and 2.5×10^4^ cells in the myeloid medium. On days 4 and 7 of differentiation, cell surface marker expression was analyzed by flow cytometry. The following fluorochrome-coupled antibodies were used for flow cytometry: CD41-PECy7 (BioLegend) clone MWReg30, CD42d-APC (eBioscience) clone 1C2, Ter119-Pacific Blue (BioLegend) clone TER-119, CD71-PerCP-Cy5.5 (BioLegend) clone RI7217, CD117-APC-H7 (BD) clone 2B8, CD11b-V500 (BD) clone M1/70 and Gr1-PerCP-Cy5.5 (BD) clone RB6-8C5. In the case of CD34^+^ HSPCs, 8×10^4^ transduced cells were seeded for megakaryocytic differentiation assays, while 5×10^4^ cells were used for combined megakaryocytic/erythroid and myeloid assays. These cells were also stained and evaluated for the expression of cell surface markers on days 7 and 10 of differentiation. The following fluorochrome-coupled antibodies were used for these assays: CD41-PECy7 (BC) clone P2, CD41-PE (BC) clone P2, CD61-PECy7 (BC) clone SZ21, CD42b-APC (BD) clone HIP1, CD235a-APC (BD) clone GA-R2, CD71-APC-AF750 (BC) clone YDJ1.2.2, CD117-PE (BC) clone 104D2D1, CD11b-APC-AF750 (BC) clone Bear1, CD14-APC (BC) clone RMO52, CD15-V500 (BD) clone HI98, CD16-PE (BC) clone 3G8 and CD66b-PECy7 (eBioscience) clone G10F5. All measurements were performed on a FACS Canto Flow Cytometer (BD Biosciences) and the data were subsequently analyzed with FlowJo v10 (FlowJo, LLC).

#### Colony-forming assays

1×10^5^ sorted CD34^+^ HSPCs were seeded in for collagen-based colony-forming assays (MegaCult™-C medium with cytokines, StemCell Technologies), as previously described (Gialesaki et al., 2018). Similarly, 10^4^ sorted CD34^+^ HSPCs were plated for methylcellulose-based colony forming assays [Human Methylcellulose Base Medium HSC002 (R&D Systems), which were supplemented with 2% streptomycin/penicillin, 10ng/mL SCF, 10ng/mL FLT3, 4U/mL EPO, 50ng/mL TPO, 50ng/mL GM-CSF, 20ng/mL IL3 and 10ng/mL IL6] according to the manufacturer’s instructions and incubated for 7 days in a 37°C incubator with 5% CO_2_ and ≥95% humidity. Serial replatings were also performed. Counting and classification of the colonies were performed with the Axiovert 40CFL microscope (Zeiss).

#### Immunophenotyping of ML-DS blasts

ML-DS blasts were analyzed for the expression of surface cell markers 5 and 14 days after lentiviral transduction. The following fluorochrome-coupled antibodies were used for the staining: CD33-PerCP-Cy5.5 (BC) clone D3HL60.251, CD117-PE (BC) clone 104D2D1, CD235a-APC(BD) clone GA-R2, CD41-PECy7 (BC) clone P2, CD42b-APC (BD) clone HIP1 and CD15-V500 (BD) clone HI98. Measurements were performed on a FACS Canto Flow Cytometer (BD Biosciences) and the acquired data were analyzed with FlowJo v10 (FlowJo, LLC).

#### Mice and bone marrow transplantations

All animal experiments were performed according to protocols approved by the local authorities (Niedersächsisches Landesamt für Verbraucherschutz und Lebensmittelsicherheit and Landesverwaltungsamt Sachsen-Anhalt). The animals were maintained under pathogen-free conditions.

#### FLC transplantation into C57BL-6J mice

Cas9 heterozygous murine FLCs were obtained by timed breeding of C57BL/6J (Jackson Laboratory) females with B6J.129(Cg)-Gt(ROSA)26^Sortm1.1(CAG-cas9*,-EGFP)^Fezh/J (Cas9 knock-in mice on C57BL/6J background) males. On embryonic day E12.5/E13.5, the fetal livers were mechanically detached from the embryos and further processed as described above. For the purposes of bone marrow transplantation, Ter119-depleted FLCs were lentivirally transduced with the pLeGO.sgGata1.4.iG2 vector and expanded *in vitro*. After at least 20 days of culture, these cells were further transduced with the pLeGO-iG2 cDNA vectors. Twenty four hours later, the FLCs were transplanted intravenously into sublethally irradiated (7.5 Gy) female C57BL/6JRj mice (aged 6-15 weeks). Secondary transplantations were performed by intravenously injecting 5×10^6^ primary bone marrow cells into sublethally irradiated (7 Gy) female C57BL/6J mice. Engraftment of the transplanted cells was evaluated by retro-orbital bleeding of the recipients every 4 weeks. When signs of disease occurred, the mice were sacrificed, and harvested bone marrow cells and splenocytes were stained with the following fluorochrome-coupled antibodies: CD3e-Pacific Blue (BioLegend) clone 17A2, B220-Pacific Blue (BioLegend) clone RA3-6B2, Lineage cocktail-Pacific Blue (BioLegend), Gr1-PerCP-Cy5.5 (BD) clone RB6-8C5, CD11b-PE (BD) clone M1/70, CD117-APC-Cy7 (BD) clone 2B8, CD117-PerCP-Cy5.5 (BD) clone 2B8, CD41-PE (BD) clone MWReg30, Sca1-PECy7 (BD) clone D7, CD34-AF647 (BD) clone RAM34, CD16/32-APC-Cy7 (BD) clone 2.4G2. All measurements were performed on a FACS Canto Flow Cytometer (BD Biosciences) and the acquired data were analyzed with FlowJo v10 (FlowJo, LLC).

#### PDX transplantation into MISTRG mice

ML-DS-patient-derived xenograft (PDX) cells were generated as previously described (Bhayadia *et al*., 2018). For xenotransplantation experiments, cells were transduced with lentiviral vectors for constitutive expression of either RUNX1A, RUNX1C and EV cDNA or sgRUNX1.1 and sgCtrl sgRNAs. 24 hours post transduction, cells were mixed in a 1:1 ratio (GFP:dTomato or GFP:E2Crimson), washed, and injected via tail vein into irradiated (2.5Gy), 8-16 week old, female MISTRG mice (Regeneron Pharmaceuticals) (Rongvaux et al., 2014), with groups of five mice per condition. A total of 1×10^6^ cells were injected per mouse. Animals were sacrificed before leukemia onset and the hematopoietic compartments were analyzed. Engraftment of the transplanted cells was evaluated via tail-vein bleeding of the recipients every 4 weeks. When signs of disease occurred, harvested bone marrow cells and splenocytes were stained with fluorochrome-coupled antibodies against human CD117 and human CD45. All measurements were performed on a FACS Canto Flow Cytometer (BD Biosciences) or a Cytoflex Flow Cytometer (BD Biosciences) and the acquired data were analyzed with FlowJo v10 (FlowJo, LLC).

#### Apoptosis assays

Apoptosis was assessed in *GATA1s-*transduced *Gata1s*-FLC overexpressing RUNX1A, CMK cells and patient-ML-DS blasts derived xenografts (PDX) after lentiviral shRNA-mediated knock down of MAX. ShRNAs against Luciferase (shCtrl) were used as controls. Apoptosis was measured using APC or FITC-conjugated Annexin V (BD Biosciences).

#### Drug responsive assay

MYC:MAX dimerization was inhibited in CMK cells, murine Gata1s-RUNX1A-FLCs, PDX blasts and CD34+ HSPCs by incubation with MYCi361 (Seleckchem) for 24h. Cell viability was accessed using the CellTiter-Glo® Cell Viability Assay (Promega) according to the manufactureŕs instructions.

#### RNA sequencing data collection and analysis

RNA samples were generated from the BM cells of diseased mice, upon sorting of live fluorescent (GFP+) cells using the FACSAria cell sorter (BD Biosciences). For control samples, sorted murine HSPCs derived from E12.5-E13.5 FLCs were used (MEPs: Lin^-^ Sca1^-^cKit^+^CD34^-^CD16/32^-^, CMPs: Lin^-^Sca1^-^cKit^+^CD34^+^CD16/32^low^, GMPs: Lin^-^Sca1^-^ cKit^+^CD34^+^CD16/32^+^, LSKs: Lin^-^Sca1^+^cKit^+^; the fluorochrome-coupled antibodies used for staining are mentioned above).

In addition, RNA samples were generated from murine FLCs transduced with SGL40C.EFS.dTomato vectors carrying a sgRNA targeting *Gata1* or a control sgRNA (sgCtrl), and with pRRL.PPT.SFFV.iGFP or doxycycline-inducible SIN40C.TRE.sfGFP.idTom.Tet3G vectors expressing RUNX1A, RUNX1C or with an empty vector control. Prior to sorting the double positive populations (dTom^+^GFP^+^) using a FACSAria cell sorter (BD Biosciences), the cells were cultured in our standard megakaryocytic differentiation medium for 72h, starting 24h post transduction. SIN40C.TRE.sfGFP.idTom.Tet3G transduced cells were expanded for up to 1 week prior to fluorescence sorting. In all cases, total RNA was extracted using the Quick-RNA Microprep Kit (Zymo Research). Paired-end libraries with 2 × 75 bp reads were prepared from the extracted RNA using the TruSeq Stranded total RNA LT Sample Prep (RiboZero Gold, Illumina) using Illumina methodology. RNA reads were aligned to GRCm38 using STAR (v2.6.0). Mapped reads were annotated using Ensembl v.91 (Aken et al., 2017; Dobin et al., 2013). Gene expression levels were quantified in fragments per kilobase of exon model per million mapped reads (FPKM). The FPKM calculation was performed using the R package edgeR including TMM normalization (Mortazavi et al., 2008; Robinson and Oshlack, 2010). Functional enrichments were calculated using gene set enrichment analysis (GSEA; v3.0) on log2-transformed FPKM values (Subramanian et al., 2005). Analysis was done based on previously described and curated gene sets and on ML-DS signatures (Schwarzer *et al*., 2017). Human gene symbols from the published leukemia gene sets were mapped to murine gene symbols using orthologue annotations provided by Ensembl considering only one-to-one orthologue relationships. Standard deviation of gene expression was calculated using log2-transformed FPKM values. Genes that showed a standard deviation >1 were selected for unsupervised hierarchical clustering.

#### Cleavage under targets and release using nuclease (CUT&RUN)

*Gata1s*-FLCs expressing doxycycline-inducible HA-RUNX1A/-RUNX1C or HA-GATA1/-GATA1s were harvested 36h and 48h after doxycline induction. Transgene expressing cells were sorted for GFP+ and dTomato+ expression. A minimum number of 100,000 cells per condition were used. CUT&RUN was performed (Skene and Henikoff, 2017)as previously described (Skene and Henikoff, 2017) using antibodies recognizing endogenous Gata1s and full length Runx1 isoforms as well as an HA-specific antibody. Sequencing was performed on an Illumina NextSeq machine at the sequencing core of the Max Planck Institute for Molecular Genetics (MPIMG, Berlin). Paired, raw reads were demultiplexed, resulting in reads 42 bases in length. The reads were quality- and adapter-trimmed using BBduk, then aligned to the mouse reference genome, (mm9), using BWA (Version: 0.7.17-r1188) in ‘aln’ mode. Mapped reads were normalized to the input cell number. For peak calling we used SEACR (version 1.1) with relaxed settings and ‘norm’ mode. Empty vector control (EV) was used as background (Meers et al., 2019a). Peaks were called using the homer software suite (v4.10) (Heinz et al., 2010). Enhancer regions were annotated by using EnhancerAtlas 2.0. We considered ESC hematopoietic progenitor, erythroblast fetal liver and erythroid fetal liver enhancers in mm9. A region was annotated as an enhancer region if it overlapped with any of the aforementioned enhancers. Overlapping peaks between transcription factors were determined with bedtools, as overlapping to any degree. Heatmaps and cluster analyses were generated using deepTools v3.4.3. (Ramirez et al., 2016). TFBS analysis was performed using the genomatix genome analyzer software suite v3.131123, TFBS Overrepresentation tool, with MatBase version 11.2, Mus musculus genome version GRCh38, ElDorado database 04-2020, and Matinspector version 8.3 for locating motif regions (Cartharius et al., 2005; Quandt et al., 1995). De novo motif discovery was performed using RSAT’s peak-motif tool (Thomas-Chollier et al., 2012) with peaks trimmed to 800 bp from the center on either side, or otherwise default settings. The discovered motifs were subsequently searched in JASPAR’s redundant vertebrate database (Fornes et al., 2020).

#### Protein isolation and Western blotting

Total cellular protein was isolated using RIPA solution (Thermo Fisher Scientific) under standard conditions. A minimum of 100,000 cells were processed for Western blotting. Protein solutions were supplemented with 4x Laemmli loading buffer (Thermo Fisher Scientific) and subsequently heated at 100°C for 10 min followed by the addition of 2-mercaptoethanol (Sigma-Aldrich). Proteins were separated by acrylamide gel electrophoresis and blotted according to standard protocols. Western blotting was performed under standard conditions using anti-RUNX1-RUNT, anti-GATA1, anti-HA, anti-Max and anti-Vinculin antibodies. Protein visualization was accomplished using anti-rabbit or anti-mouse HRP-conjugated secondary antibodies (Abcam) and Amersham™ ECL Prime Western Blotting Detection Reagent (Thermo Fisher Scientific) according to the manufacturer’s instructions, followed by detection of chemiluminescent signals on a ChemiDoc station (BioRad).

#### Co-immunoprecipitation

A minimum of 1×10^7^ K562 cells (Western blotting) or 1×10^8^ CMK cells (Mass spectrometry) stably transduced with doxycycline-inducible 1xHA-tagged RUNX1 or GATA1 isoforms as well as the empty control (EV) were induced for transgene expression by application of 0.25 - 1μg/ml doxycycline. 36-48h after induction, cells were gently lysed for 30 minutes on ice in 3 cell volumes of hypotonic lysis buffer (10 mM Tris/HCl (pH 7.4); 2 mM MgCl2; 0.5% TritonX) supplemented with PIC (Sigma-Aldrich) and PMSF protease inhibitor (Thermo Fisher Scientific), followed by solubilization of nuclei using a cell homogenisator. Nuclei were precipitated by centrifugation at 8500 rpm for 15 minutes at 4°C. Nuclear proteins were isolated by mechanistic disruption of the nuclear pellet by dounce-homogenization and incubation in nuclear extraction buffer (20 mM HEPES; 10 mM KCl; 1 mM EDTA; 350 mM NaCl; 20% glycerol supplemented with (Sigma-Aldrich), PMSF protease inhibitor (Thermo Fisher Scientific) and sodium-orthovanadate (Merckmillipore) for 30 minutes on ice. Insoluble cellular components were precipitated by centrifugation for 30 minutes at 13,000 rpm and 4°C. Protein complexes were immunoprecipitated using anti-HA-conjugated magnetic beads (Pierce^TM^ anti-HA magnetic beads; Thermo Fischer Scientic) following the manufactureŕs protocol. Protein complexes were eluted from the beads via boiling for 10 minutes (Western blotting) or incubation at 37°C in 8M urea solution supplemented with 10 mM DTT (Carl Roth) for 30 minutes and processed for mass spectrometry.

#### Mass spectrometric analysis of protein complexes

Protein complexes were proteolyzed with trypsin following the FASP (filter aided sample preparation) protocol (Wisniewski et al., 2009). Samples were analyzed by LC/MS/MS using 180-min gradients on an U3000 nano-HPLC system coupled to a Q-Exactive Plus or an Orbitrap Fusion mass spectrometer (both Thermo Fisher Scientific). Peptides were separated on reversed phase C18 columns (trapping column: Acclaim PepMap 100, 300 μm × 5 mm, 5μm, 100 Å (Thermo Fisher Scientific); separation column: µPAC 50 cm C18 (Pharmafluidics) or self-packed RP C18 emitter column (PicoFrit, 75 μM × 250– 500 mm, 15 µm tip diameter (New Objective), packed with *ReproSil-Pur* C18*-*AQ, 1.9 μm, 120 Å, Dr. Maisch, Germany). After desalting the samples on the trapping column, peptides were eluted and separated using linear gradients from 3% to 40% B (solvent A: 0.1% (v/v) formic acid in water, solvent B: 0.08% (v/v) formic acid in acetonitrile) with a constant flow rate of 300 nl/min over 180 or 240 min. For data acquisition with the Orbitrap Fusion MS, a data-dependent top 5s method was applied. FTMS survey scans were performed in the *m/z* range 350-1500 (R = 120,000 at *m/z* 200, target of the automatic gain control (AGC) 400,000, maximum injection time (IT) 50 ms). MS/MS scans of the most abundant signals of the survey scans were acquired in parallel by FTMS (HCD (higher energy collision-induced dissociation) at 28% NCE (normalized collision energy), AGC 50,000, IT 120 ms) and CID in linear ion trap (collision induced dissociation), 35% NCE, AGC 10,000, IT 35 ms), isolation window was 1.5 Th. For MS data acquisition with the Q-Exactive Plus mass spectrometer, high-resolution full scans (m/z 375 to 1799, R = 140,000 at m/z 200) in the orbitrap were followed by high-resolution product ion scans (R = 17,500) of the 10 most intense signals in the full-scan mass spectrum (isolation window 2 Th); AGC was set to 3 x 10^6^ (MS) and 200,000 (MS/MS), IT were set to 100 ms (MS) and 150 ms (MS/MS). For acquisitions on both instruments, precursor ions with charge states <2+ and >6+ or were excluded from fragmentation. Dynamic exclusion was enabled (duration 60 seconds, window 3 ppm).

Raw data were processed using Proteome Discoverer 2.4 (Thermo Fisher Scientific). MS/MS data were searched against the UniProt database (version Nov. 2019, tax. *homo sapiens*, 73801 entries) using SequestHT. Label-free quantification of proteins was based on extracted peak areas of corresponding peptide precursor ions, quantitative data were normalized on total protein abundance.

Proteins were selected on a minimum of 3 unique measured peptides. Common contaminates (MaxQuant contaminates.fasta, MPI Martinsried) were excluded. Abundance log2 ratios of either RUNX1A or RUNX1C versus empty control of greater than 1 (p-value < 0.05) were assumed as significantly bound. Pathway enrichment analysis was done for either unique or common bound proteins for RUNX1A and RUNX1C using STRING pathway enrichment analysis tool as plugin of Cytoscape software (Doncheva et al., 2019; Shannon et al., 2003). Cofactor core complexes were identified using the Corum database (Giurgiu et al., 2019).

#### Statistical analysis

Statistical evaluation was performed using Student’s t-tests, Mann-Whitney tests, and one-way or two-way ANOVA. The Kaplan-Meier method and log-rank tests were used to estimate overall survival and to compare differences between survival curves, respectively. All data are presented as mean ± standard deviation (SD). Calculations were performed using GraphPad Prism 6/7 (STATCON). All statistical tests and sample numbers are disclosed in the respective figure legends/supplementary tables.

## Supplementary Figures

**Figure S1.** CRISPR-Cas9 screen reveals *RUNX1* dependency in ML-DS (related to Figure 1)

**Figure S2.** Increased RUNX1A:RUNX1C ratio induces malignant ML-DS phenotype (related to Figure 2)

**Figure S3.** *RUNX1A* synergizes with *Gata1s* in leukemic transformation of murine fetal liver cells (related to Figure 3)

**Figure S4.** Distinct RUNX1A or RUNX1C protein interaction networks and their effects on gene regulation (related to Figure 4 and 5)

**Figure S5.** RUNX1A exerts oncogenic effects via MYC:MAX (related to Figure 5)

## Supplementary Tables

**Table S1.** Hsa21 CRISPR/Cas9 sgRNA library and screen results (related to Figure 1)

**Table S2.** Patient characteristics (related to Figure 1-2 and 5, Figure S1-2 and S5)

**Table S3.** GSEA of sgGata1s- or sgCtrl-FLCs transduced with *RUNX1A*, *RUNX1C* or EV (related to Figure 3)

**Table S4.** GSEA of murine ML-DS leukemia samples compared to HSPCs (related to Figure 3)

**Table S5.** Transcription factor overrepresentation analysis of CUT&RUN peaks (related to Figure 5)

