## Supplementary Information for "*RUNX1* isoform disequilibrium in the development of trisomy 21 associated myeloid leukemia"

### Supplementary figures

**Figure S1.** CRISPR-Cas9 screen reveals *RUNX1* dependency in ML-DS (related to Figure 1)

**Figure S2.** Increased RUNX1A:RUNX1C ratio induces malignant ML-DS phenotype (related to Figure 2)

**Figure S3.** *RUNX1A* synergizes with *Gata1s* in leukemic transformation of murine fetal liver cells (related to Figure 3)

**Figure S4.** Distinct RUNX1A or RUNX1C protein interaction networks and their effects on gene regulation (related to Figure 4 and 5)

**Figure S5.** RUNX1A exerts oncogenic effects via MYC:MAX (related to Figure 5)

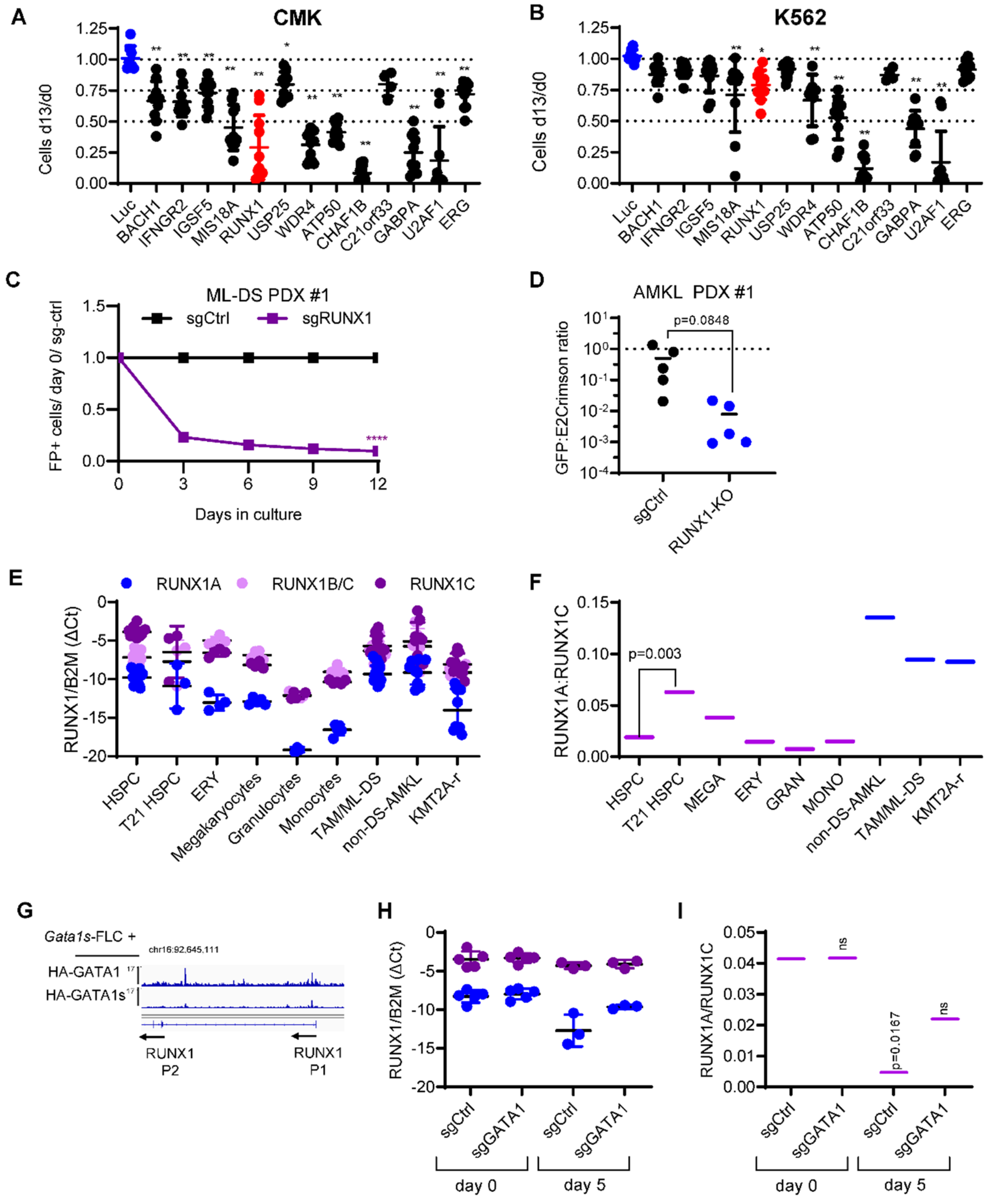

**Figure S1. CRISPR-Cas9 screen reveals *RUNX1* dependency in ML-DS (related to Figure 1)**
(A-B) Dot plots showing the number of sgRNA-transduced (A) CMK and (B) K562 cells (with stable Cas9 expression) after 13 days of culture normalized to day 0 (n=3 per sgRNA, one-way ANOVA). $P_{ANOVA}$<0.05, **$P_{ANOVA}$<0.01.
(C) Percentage of GFP$^+$ sgRNA-transduced (sgCtrl and sgRUNX1.1) ML-DS PDX cells normalized to day 0 and sgCtrl (mean±s.d., n=4, two-way ANOVA, **** $P_{ANOVA}$ <0.0001).
(D) Dot plot showing the number of sgRNA-transduced (sgCtrl and sgRUNX1.1) AMKL PDX cells (with stable Cas9 expression) after 12 days of culture normalized to day 0 (n=5, two-tailed unpaired t-test).
(E) Expression of *RUNX1A, RUNX1B/C* and *RUNX1C* isoforms normalized to the expression of β2-microglobulin (*B2M*) in CD34$^+$ HSPCs, erythrocytes, megakaryocytes, granulocytes and monocytes isolated from healthy donors, as well as in leukemic blasts from TAM/ML-DS, non-DS-AMKL and KMT2A-rearraged patients.

(F) Ratio of *RUNX1A* to *RUNX1C* expression in CD34⁺ HSPCs, erythrocytes, megakaryocytes, granulocytes and monocytes isolated from healthy donors, as well as in leukemic blasts from TAM/ML-DS, non-DS-AMKL and KMT2A-rearraged patients.
(G) IGV snapshots of the *RUNX1* gene locus, showing occupancy of HA-GATA1 and HA-GATA1s in *Gata1s*-FLCs after doxycycline induced HA-GATA1/GATA1s expression or in EV control expressing cells. The *RUNX1* promoters are indicated with P1 and P2. The tracks display coverage (RPKM) (left). Scale and chromosome location are shown (top).
(H) Expression of *RUNX1A* and *RUNX1C* isoforms normalized to the expression of β2-microglobulin (*B2M*) in cord blood (CB) HSPCs expressing Cas9 and sgRNAs against *GATA1* (sgGATA1) or control sgRNAs (sgCtrl) at days 0 and 5 after transduction. Data from at least 3 replicates are shown.
(I) Ratio of *RUNX1A* to *RUNX1C* expression in neonatal MEPs expressing sgRNAs against *GATA1* (sgGATA1) or control sgRNAs (sgCtrl) at days 0 and 5 after transduction (n≥3, one-way ANOVA compared to sgCtrl at day 0).

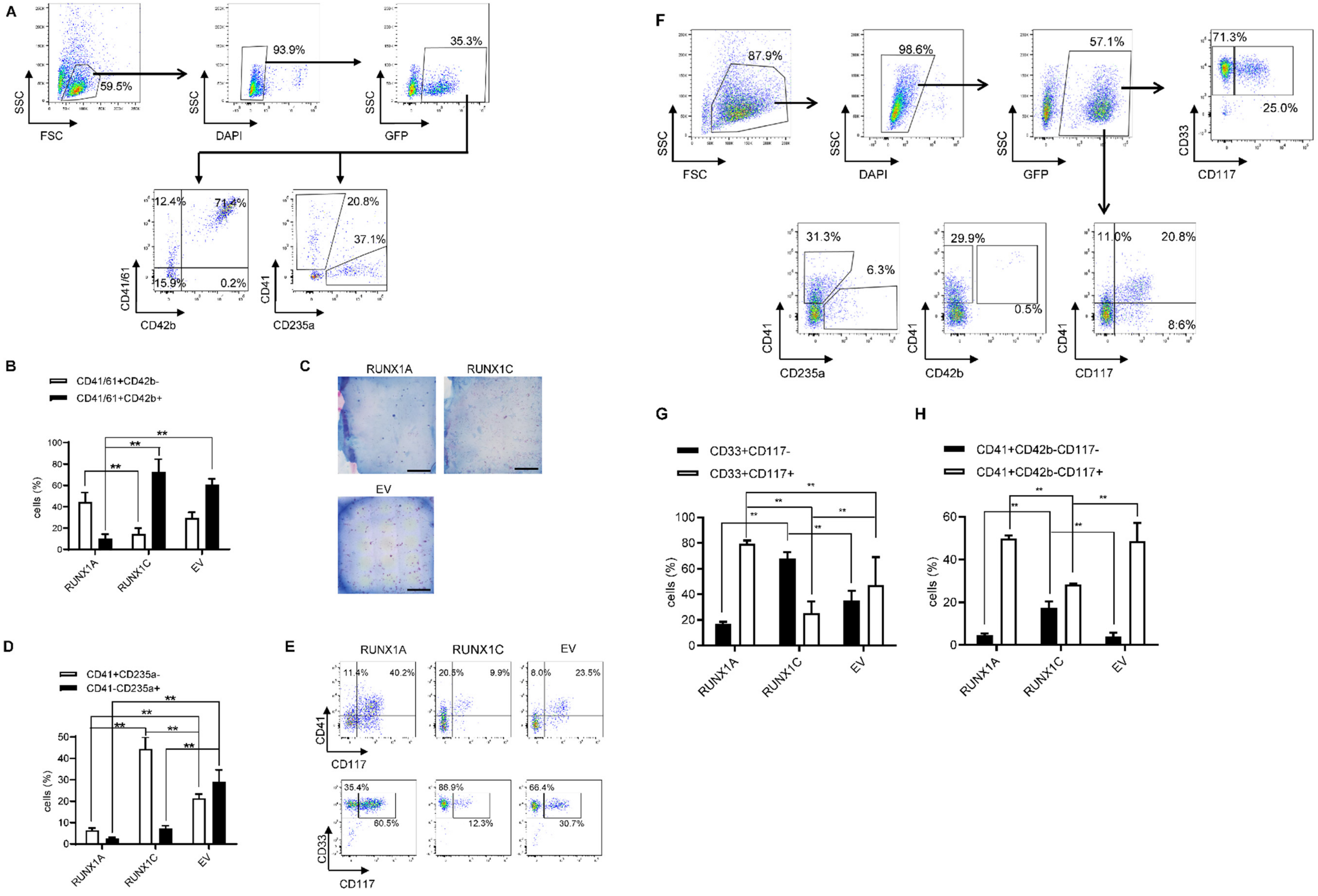
A
59.5%
93.9%
35.3%
SSC
FSC
DAPI
GFP
12.4%
71.4%
15.9%
0.2%
CD41/61
CD42b
20.8%
37.1%
CD41
CD235a
B
CD41/61+CD42b-
CD41/61+CD42b+
cells (%)
RUNX1A
RUNX1C
EV
C
RUNX1A
RUNX1C
EV
D
CD41+CD235a-
CD41-CD235a+
cells (%)
E
RUNX1A
RUNX1C
EV
11.4%
40.2%
20.5%
9.9%
8.0%
23.5%
CD41
CD117
35.4%
60.5%
86.9%
12.3%
66.4%
30.7%
CD33
CD117
F
87.9%
98.6%
57.1%
71.3%
25.0%
SSC
FSC
DAPI
GFP
CD33
CD117
31.3%
6.3%
29.9%
0.5%
11.0%
20.8%
8.6%
CD41
CD235a
CD42b
CD117
G
CD33+CD117-
CD33+CD117+
cells (%)
H
CD41+CD42b-CD117-
CD41+CD42b-CD117+
cells (%)

**Figure S2. Increased RUNX1A:RUNX1C ratio induces malignant ML-DS phenotype (related to Figure 2)**

(A-E) Neonatal $CD34^+$ HSPCs were lentivirally transduced with *RUNX1A*-, *RUNX1C*- or control (empty) overexpressing vectors.

(A) Gating strategy applied in flow cytometric analysis of transduced $CD34^+$ HSPCs. Data from the control (empty vector; EV) cells are shown. Cells were first gated on the forward scatter (FSC)/side scatter (SSC) plot (upper left). Live cells (upper middle) were gated to detect $GFP^+$ cells (upper right). These cells were further gated to detect differentiation-specific markers (CD41/61/CD42b, bottom left; CD41/CD235a, bottom right).

(B) Percentage of transduced neonatal HSPCs expressing CD41/CD61/CD42b cell surface markers on day 10 of megakaryocytic differentiation. Data from three independent replicates are shown as mean±s.d. Two-way ANOVA. $^{**}P_{ANOVA}<0.01$

(C) Photographs of CFU-Mk assays (scale bar 5mm).

(D) Percentage of transduced neonatal HSPCs expressing CD41/CD235a cell surface markers on day 10 of erythroid-megakaryocytic differentiation. Data from three independent replicates are shown as mean±s.d. Two-way ANOVA. $^{**}P_{ANOVA}<0.01$

(E) Flow cytometric analysis of CD41/CD117 (upper panel) and CD33/CD117 surface marker expression on day 10 of erythroid-megakaryocytic differentiation. Representative dot plots are shown. The percentage of cells belonging to each immunophenotype is indicated in the corresponding gate.population.

(F) Gating strategy applied in flow cytometric analysis of transduced ML-DS blasts. Data from the control (empty vector; EV) cells are shown. Cells were first gated on the forward scatter (FSC)/side scatter (SSC) plot (upper far left). Live cells (upper middle left) were gated to detect $GFP^+$ cells (upper right). These cells were further gated to detect differentiation-specific markers (CD33/CD117, upper far right; CD41/CD235a, bottom left; CD41/CD42b, bottom middle; CD41/CD117, bottom right).

(G-H) Percentage of transduced ML-DS blasts expressing the CD33/CD117 (G) and CD41/CD117 (H) cell surface markers, 14 days post-transduction. Data from two replicates are shown as mean±s.d. Two-way ANOVA. $^{**}P_{ANOVA}<0.01$

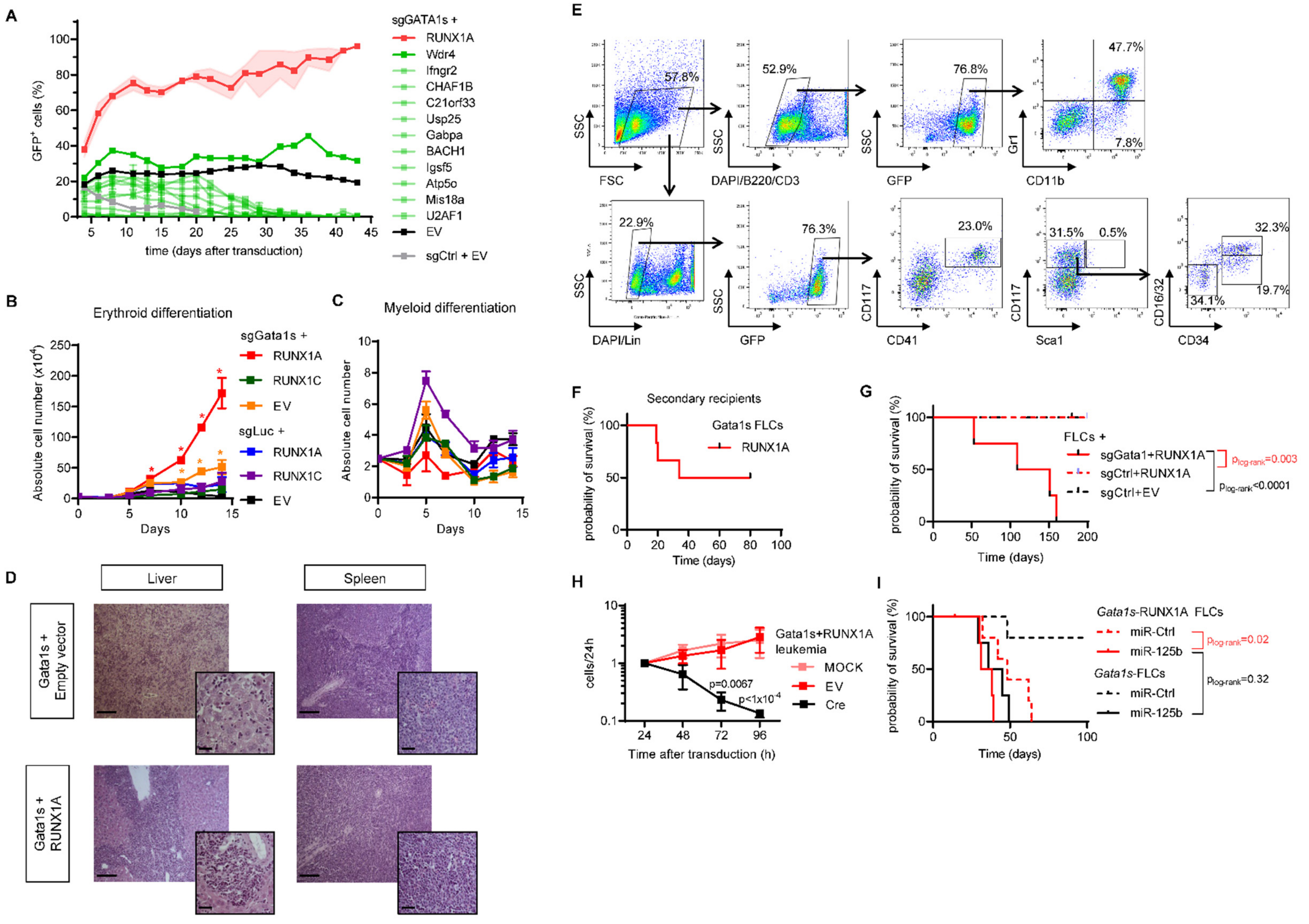
A
sgGATA1s +
RUNX1A
Wdr4
Ifngr2
CHAF1B
C21orf33
Usp25
Gabpa
BACH1
Igsf5
Atp5o
Mis18a
U2AF1
EV
sgCtrl + EV
GFP+ cells (%)
time (days after transduction)
B
Erythroid differentiation
sgGata1s +
RUNX1A
RUNX1C
EV
sgLuc +
RUNX1A
RUNX1C
EV
Absolute cell number (x10^4)
Days
C
Myeloid differentiation
Absolute cell number
Days
D
Liver
Spleen
Gata1s + Empty vector
Gata1s + RUNX1A
E
57.8%
52.9%
76.8%
47.7%
7.8%
SSC
FSC
DAPI/B220/CD3
GFP
Gr1
CD11b
22.9%
76.3%
23.0%
31.5%
0.5%
32.3%
19.7%
34.1%
DAPI/Lin
CD117
CD41
Sca1
CD16/32
CD34
F
Secondary recipients
Gata1s FLCs
RUNX1A
probability of survival (%)
Time (days)
G
FLCs +
sgGata1+RUNX1A
sgCtrl+RUNX1A
sgCtrl+EV
plog-rank=0.003
plog-rank<0.0001
probability of survival (%)
Time (days)
H
Gata1s+RUNX1A leukemia
MOCK
EV
Cre
p=0.0067
p<1x10-4
cells/24h
Time after transduction (h)
I
Gata1s-RUNX1A FLCs
miR-Ctrl
miR-125b
Gata1s-FLCs
miR-Ctrl
miR-125b
plog-rank=0.02
plog-rank=0.32
probability of survival (%)
Time (days)

**Figure S3. RUNX1A synergizes with Gata1s in leukemic transformation of murine fetal liver cells (related to Figure 3)**

Cas9-knock-in Ter119$^-$ FLCs were lentivirally transduced with *Gata1* (sgGata1s) or control (sgCtrl) sgRNAs, as well as with the indicated cDNA or empty vector control (EV).

(A) Growth curves after transduction of cDNAs of select candidate genes from the sgRNA drop out screen. Percentage of fluorescent cells in cDNA transduced samples. Data from 4 replicates are shown as mean±s.d.

(B-C) Number of fluorescent cells after transduction with *RUNX1A*-, *RUNX1C*- or control (EV) vectors and culture in media promoting erythroid (B) or myeloid (C) differentiation. Data from 2 replicates are shown as mean±s.d. Two-way ANOVA. *$P_{ANOVA}$<0.05, **$P_{ANOVA}$<0.01.

(D) Representative microscopic images of liver and spleen sections from the indicated experimental groups (2 µm sections, H&E stained, 100-fold magnified, scale bar 500µm, highlighted segment 400-fold magnified, scale bar 80µm).

(E) Gating strategy applied in flow cytometric analysis of transduced FLCs isolated from the bone marrow of diseased mice. Data from the control (empty vector; EV) cells of healthy recipients are shown. Cells were first gated on the forward scatter (FSC)/side scatter (SSC) plot (upper far left). Live cells, negative for lymphoid lineage (B220/CD3) (upper middle left) were gated to detect GFP$^+$ cells (upper right). These cells were further gated to detect myeloid-specific markers (CD11b/Gr1, upper right). Live cells, negative for mature myeloid lineages (Lin, lower far left) were gated to detect GFP$^+$ cells (lower left). Cells were further gated for CD41/CD117 (lower middle), Sca1/CD117 (lower right) CD16/32/CD34 (bottom right).

(F) Kaplan-Meier survival curve of secondary recipients transplanted with BM-derived leukemic cells from 3 primary recipients.

(G) Kaplan-Meier survival curve of mice transplanted with either FLCs transduced with sgCtrl+empty vector (EV; n=11), sgCtrl+RUNX1A (n=5), sgGata1+EV (n=12) or sgGata1+RUNX1A (n=4; log-rank test).

(H) Number of BM leukemic cells derived from diseased mice (normalized to 24h) after transduction with a Cre recombinase-expressing vector. Data from three replicates are shown as mean±s.d. Two-way ANOVA.

(I) Kaplan-Meier survival curve of mice transplanted with either *Gata1s*-FLCs or *RUNX1A*-*Gata1s*-FLCs expressing miR-125b or control (miR-Ctrl) (n=4-5; log-rank test).

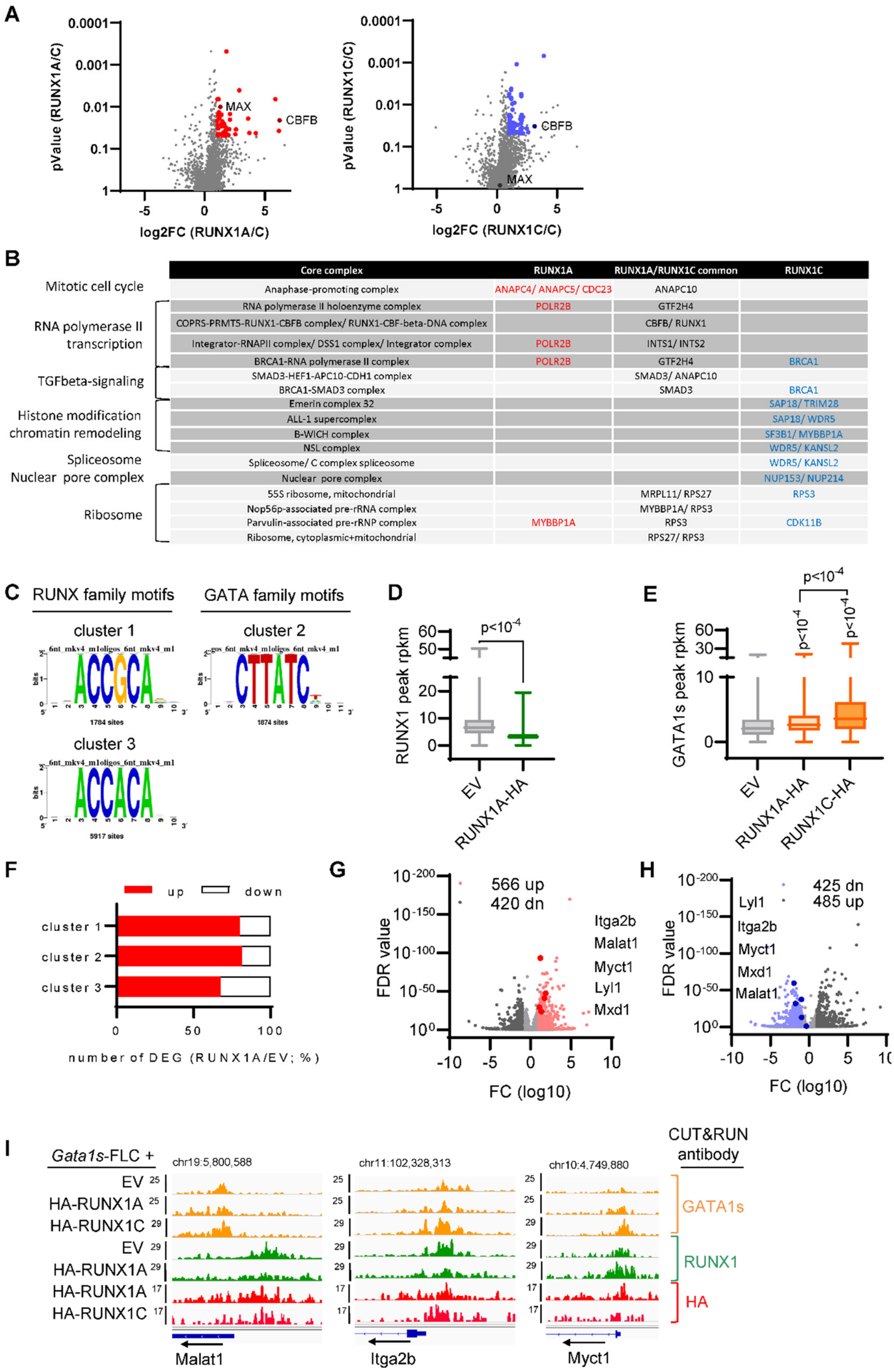


**Figure S4. Distinct RUNX1A or RUNX1C protein interaction networks and their effects on gene regulation (related to Figure 4 and 5)**

(A) Volcano plots showing enriched proteins after co-immunoprecipitation (Co-IP) of either HA-tagged RUNX1A or RUNX1C. CMK cells were induced with doxycycline for 36h before Co-IP. Proteins with a log fold change of 1 and a p-value of 0.05 in RUNX1A (red) or RUNX1C (blue) compared to the empty control were deemed significantly bound. Known RUNX1 cofactor CBFbeta and Myc-association factor X (MAX) are indicated by name.

(B) Protein core complexes of RUNX1A (red) or RUNX1C (blue) co-bound proteins. The table shows complexes for selected co-bound proteins. Data were generated using the Corum database (Giurgiu et al., 2019).

(C-I) CUT&RUN analyses of Cas9-knock-in Ter119$^-$ FLCs lentivirally transduced with *Gata1*-targeting sgRNAs, as well as with *HA-RUNX1A, HA-RUNX1C* or control (EV) vector.

(C) De novo motif discovery using sequences below RUNX1 peaks in CUT&RUN clusters (1-3). The most abundant motif in each cluster is shown. The number of the respective motif sites within each cluster are shown below.

(D) RUNX1 binding intensities (peak RPKM) at regulatory regions (enhancer and promoter) in RUNX1A-HA or empty vector control (EV) cells. (Two-way ANOVA).

(E) GATA1s binding intensities (peak RPKM) at regulatory regions (enhancer and promoter) in RUNX1A-HA, RUNX1C-HA or empty vector control (EV) cells. (Two-way ANOVA).

(F) Number of up- and downregulated genes within each CUT&RUN cluster upon doxycycline-induced *HA-RUNX1A* expression, normalized to the empty vector (EV) control (logFC >1; FDR < 0.1). Graphs depict the percentage of up- and downregulated genes normalized to the total number of deregulated genes per cluster.

(G) Volcano plot showing up- and downregulated genes upon doxycycline-induced *HA-RUNX1A* expression in *Gata1s*-FLCs. Upregulated genes are shown as light red dots. Representative genes with MYC:MAX or MAX binding motifs in their promoter regions are shown in dark red. Gene names are listed to the right according to their FDR values.

(H) Volcano plot showing up- and downregulated genes upon doxycycline-induced *HA-RUNX1C* expression in *Gata1s*-FLCs. Upregulated genes are shown as light blue dots. Representative genes with Myc:Max or Max binding motifs in their promoter regions are shown in dark blue. Gene names are listed to the left according to their FDR values.

(I) IGV snapshots of the *Malat1*, *Itga2b* and *Myct1* gene promoters, showing occupancy of endogenous GATA1s, RUNX1 and HA-RUNX1A/RUNX1C in *Gata1s*-FLCs after doxycycline induced HA-RUNX1A/RUNX1C expression or in EV control expressing cells. The tracks display coverage (RPKM) (left). Scale and chromosome location are shown (top).

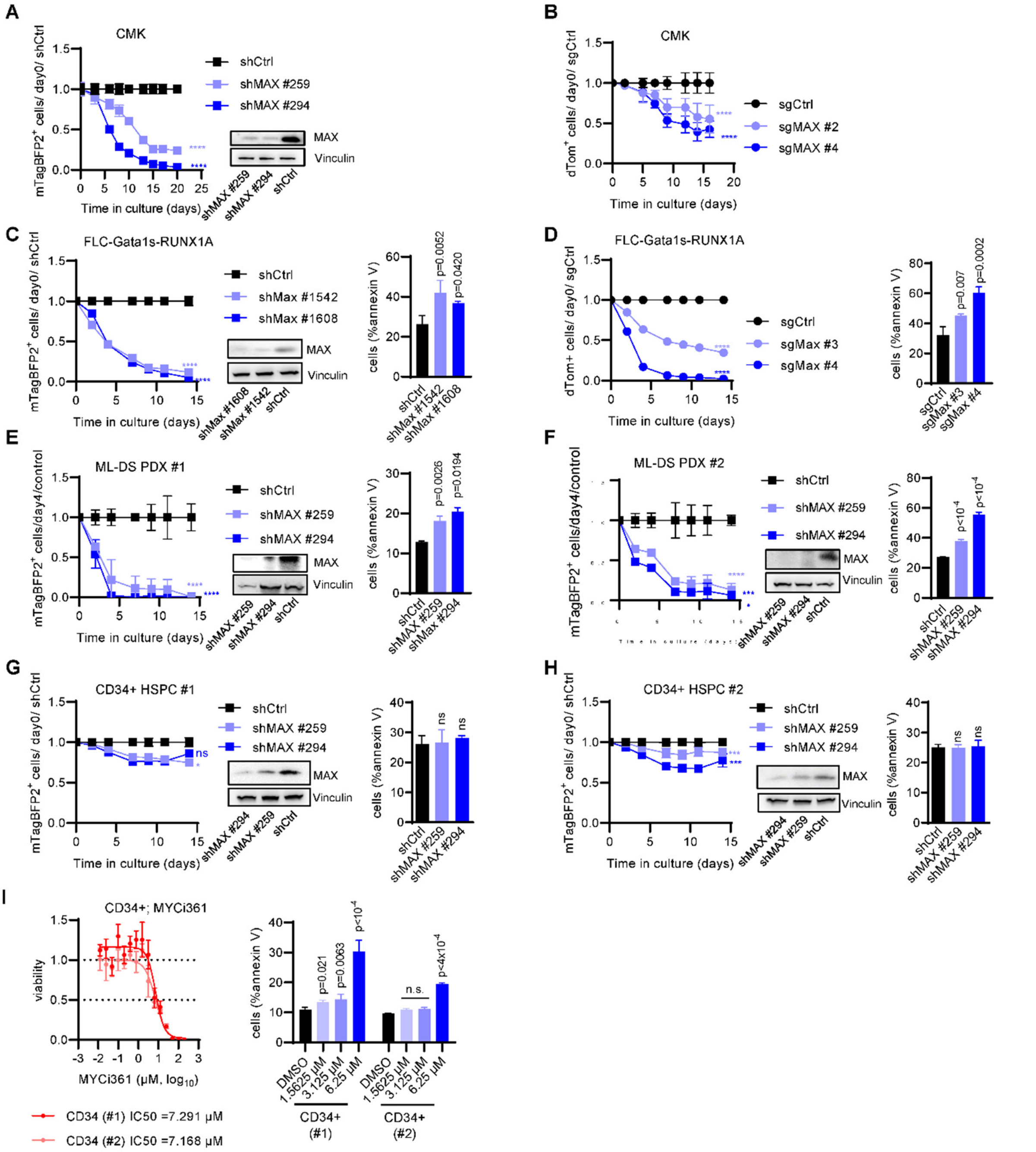


**Figure S5. RUNX1A exerts oncogenic effects via MYC:MAX (related to Figure 5)**

(A) Percentage of shMAX ($BFP2^+$) transduced CMK cells normalized to shCtrl and to day 0 (mean±s.d., n=3, one-way ANOVA) (left). Representative Western blot confirming shRNA-mediated knockdown of MAX using anti-MAX and anti-Vinculin (loading control) antibodies.

(B) Percentage of sgMAX ($dTomato^+$) transduced CMK cells normalized to sgCtrl and to day 0 (mean±s.d., n=3, one-way ANOVA).

(C) Percentage of shMax ($BFP2^+$) transduced *RUNX1A-Gata1s*-FLCs cells normalized to shCtrl and to day 0 (mean±s.d., n=3, one-way ANOVA). Representative Western blot confirming shRNA-mediated knockdown of Max using anti-Max and anti-Vinculin (loading control) antibodies. Bar graph showing percentage of $AnnexinV^+$ FLC-*Gata1s*-RUNX1A cells after shRNA-mediated knockdown of Max at day 2 (mean ± SD; n=3; one-way ANOVA).

(D) Percentage of sgMax ($dTomato^+$) transduced *RUNX1A-Gata1s*-FLCs cells normalized to sgCtrl and to day 0 (mean±s.d., n=3, one-way ANOVA). Bar graph showing percentage of $AnnexinV^+$ FLC-*Gata1s*-RUNX1A cells after sgRNA-mediated knockout of Max at day 2 (mean ± SD; n=3; one-way ANOVA).

(E-F) Percentage of shMAX (BFP2$^+$) transduced ML-DS PDX cells (PDX #1; PDX #2) normalized to shCtrl and to day 0 (mean±s.d., n=3, one-way ANOVA). Representative Western blot confirming shRNA-mediated knockdown of Max using anti-MAX and anti-Vinculin (loading control) antibodies. Bar graph showing percentage AnnexinV$^+$ ML-DS PDX cells (PDX #1; PDX #2) after shRNA-mediated knockdown of MAX at day 2 (mean ± SD; n=3; one-way ANOVA).
(G-H) Percentage of shMAX (BFP2$^+$) transduced CD34$^+$ HSPCs cells (#1 and #2) normalized to shCtrl and to day 0 (mean±s.d., n=3, one-way ANOVA). Representative Western blot confirming shRNA-mediated knockdown of MAX using anti-MAX and anti-Vinculin (loading control) antibodies. Bar graph showing percentage of AnnexinV$^+$ CD34$^+$ HSPCs cells after shRNA-mediated knockdown of MAX at day 2 (mean±s.d.; n=3; one-way ANOVA).
(I) Dose response curves for the MYC:MAX dimerization inhibitor MYCi361 in CD34$^+$ HSPCs 24h after treatment *in vitro* (left). The IC50-values are depicted below the graph. Bar graph showing the percentage of AnnexinV$^+$ cells after treatment with the indicated doses of MYCi361 in comparison to the DMSO control (right; mean±s.d.; n=3; one-way ANOVA).

**Supplementary Tables**

**Table S1.** Hsa21 CRISPR/Cas9 sgRNA library and screen results (related to Figure 1)

**Table S2.** Patient characteristics (related to Figure 1-2 and 5, Figure S1-2 and S5)

**Table S3.** GSEA of sgGata1s- or sgCtrl-FLCs transduced with *RUNX1A*, *RUNX1C* or EV (related to Figure 2)

**Table S4.** GSEA of murine ML-DS leukemia samples compared to HSPCs (related to Figure 3)

**Table S5.** Transcription factor overrepresentation analysis of CUT&RUN peaks (related to Figure 5)

**Table S2.** Patient characteristics (related to Figure 1-2 and 5, Figure S1-2 and S5)

| | gender | age at diagnosis (years) | WBC ($x10^9/L$) | hemoglobin (g/dl) | BM blasts (%) | CNS | SCT | molecular genetics | Cytogenetics (karyotype) | response | relapse | Related to |
|---|---|---|---|---|---|---|---|---|---|---|---|---|
| **ML-DS PDX#1** | m | 2 1/6 | 32.5 | 7.9 | 70.5 | no | yes | GATA1 mutation | 47,XY,t(3;13)(q?26;q?13~14)del(13)(q?14q22), +21c[cp14]/47,sl,del?(15)(q?)[cp2]/46,XY[1] | NR, CCR:after SCT | no | Figure 1; 2; 5; S1; S5 |
| **ML-DS PDX#2** | f | 1 1/12 | 168 | 7.1 | 16 | no | no | NRAS mutation | ns | CCR | no | Figure 5;2; S5 |
| **ML-DS PDX#3** | m | 1 1/3 | 4.8 | 12 | 10.5 | no | no | GATA1 mutation | ns | CCR | no | Figure 5 |
| **AMKL PDX#1** | m | 1 | 40 | 9.9 | 64 | no | yes | KMT2A mutation | 46,XY[15].nuc ish 3q26(EVI1x2)[100/100], 8q22(RUNX1T1x2),21q22(RUNX1x2)[98/100], 11q23(MLLx2)[99/100],16q22(CBFBx2)[100/100] 17q21.1(RARAx2)[100/100] | CCR | yes | Figure S1 |
| **TAM** | f | 0 | 107 | 13.1 | 86 (PB) | ns | no | GATA1 mutation | ns | CCR | ML-DS | Figure 1 |

ns = not specified; BM = bone marrow; PB = peripheral blood, WBC = white blood cell count; NR = non-response; CCR complete continuous remission; SCT = stem cell transplantation
